# Disruption of origin chromatin structure by helicase activation in the absence of DNA replication

**DOI:** 10.1101/2021.03.17.435814

**Authors:** Rachel A. Hoffman, David M. MacAlpine

## Abstract

Prior to initiation of DNA replication, the eukaryotic helicase, Mcm2-7, must be activated to unwind DNA at replication start sites in early S-phase. To study helicase activation within origin chromatin, we constructed a conditional mutant of the polymerase α subunit Cdc17 (or Pol1) to prevent priming and block replication. Recovery of these cells at permissive conditions resulted in the generation of unreplicated gaps at origins, likely due to helicase activation prior to replication initiation. We used micrococcal nuclease (MNase)-based chromatin occupancy profiling under restrictive conditions to study chromatin dynamics associated with helicase activation. Helicase activation in the absence of DNA replication resulted in the disruption and disorganization of chromatin which extends up to one kilobase from early, efficient replication origins. The CMG holo-helicase complex also moves the same distance out from the origin, producing single-stranded DNA that activates the intra-S-phase checkpoint. Loss of the checkpoint did not regulate the progression and stalling of the CMG complex, but rather resulted in the disruption of chromatin at both early and late origins. Finally, we found that the local sequence context regulates helicase progression in the absence of DNA replication, suggesting that the helicase is intrinsically less processive when uncoupled from replication.

## Introduction

The stable DNA double helix must be melted and unwound into two single strands prior to the initiation of DNA replication. In eukaryotic organisms, the Mcm2-7 complex serves as the replicative helicase responsible for unwinding the duplex DNA at the origin and for continued separation of strands ahead of the replisome at the replication fork. Mcm2-7 is loaded at replication start sites during G1 phase of the cell cycle and is made fully active by the association of Cdc45 and the GINS complex to form the CMG (Cdc45-Mcm2-7-GINS) holo-helicase in early S-phase (Bell and Labib 2016). While genetic and biochemical studies have begun to elucidate the factors that are necessary and sufficient for helicase activation, we have little understanding of how the active helicase initially melts the duplex DNA in the context of the local chromatin environment or of the mechanisms that regulate CMG progression through chromatin.

Prior to helicase activation, replication origins must be selected and licensed. The conserved origin recognition complex (ORC) selects potential start sites (Bell and Stillman 1992). During G1 phase of the cell cycle, at least two copies of Mcm2-7 are recruited to each origin and loaded by ORC, Cdt1, and Cdc6 (Tanaka and Diffley 2002; Randell et al. 2006; Remus et al. 2009). These Mcm2-7 complexes encircle the DNA helix and interact to form double hexamers (Remus et al. 2009). In early S-phase, DDK and S-CDK phosphorylation recruit various factors to the helicase including Cdc45, polymerase ε, and the GINS complex; together, these form the pre-initiation complex (Tanaka et al. 2007; Zegerman and Diffley 2010; Heller et al. 2011). At this point, the Mcm2-7 double hexamer separates into two independent CMG complexes (Miyazawa-Onami et al. 2017; Douglas et al. 2018). Mcm10 association is necessary to activate the CMG holo-helicase to initially unwind DNA (Kanke et al. 2012; Van Deursen et al. 2012; Watase et al. 2012). Now active, the CMGs must unwind the DNA double helix and pass each other on single-stranded DNA (ssDNA) in order for Mcm2-7 to travel with the correct orientation on the leading strand (Georgescu et al. 2017; Douglas et al. 2018). Following origin unwinding, replication begins on both the leading and lagging strands with the synthesis of primers by polymerase α, which are extended by polymerases δ and ε (Bell and Labib 2016).

The local chromatin environment at replication origins is important for origin selection and helicase loading (Gutiérrez and MacAlpine 2016). In budding yeast, ORC binds to a T-rich sequence (ACS) (Broach et al. 1983; Bell and Stillman 1992) that is necessary, but not sufficient for origin function (Breier et al. 2004). In addition to the primary sequence, the local chromatin structure is also thought to contribute to origin activity (Thoma et al. 1984; Eaton et al. 2010; Berbenetz et al. 2010). In higher eukaryotes, open chromatin organization serves as the major determinant of origin identity. (MacAlpine et al. 2010; Miotto et al. 2016; Long et al. 2020). Experiments at the yeast origin ARS1 initially established that specific nucleosome positioning around ARS1 is necessary for origin function (Simpson 1990). ORC binding establishes a precise positioning for the nucleosomes flanking origins across the budding yeast genome (Eaton et al. 2010; Berbenetz et al. 2010). During G1 phase, a Cdc6-dependent shift in the flanking nucleosome occurs around the origin, suggesting that Mcm2-7 loading expands the nucleosome-free region (NFR) at replication start sites (Belsky et al. 2015). Though the minimal set of replication and chromatin factors necessary for origin activation have been identified on chromatinized templates *in vitro* (Kurat et al. 2017; Azmi et al. 2017; Devbhandari et al. 2017), the chromatin dynamics in early S phase that accompany pre-initiation complex formation and helicase activation remain unaddressed.

Prior to the start of replication elongation, helicase activation and subsequent DNA unwinding in early S phase serve as a model for the uncoupling of helicase and polymerase activity. In eukaryotes, helicase-polymerase uncoupling was initially observed in *Xenopus* extracts when aphidicolin treatment to inhibit polymerases α and δ caused extensive negative supercoiling, indicating that DNA could be unwound by the helicase even in the absence of replication (Walter and Newport 2000; Pacek and Walter 2004). Subsequently, uncoupling of the helicase and at least leading strand polymerization have been observed in multiple contexts that disrupt replication, including polymerase inhibition, DNA damage, and dNTP depletion (Ercilla et al. 2020; Nedelcheva et al. 2005; Sogo et al. 2002; Gan et al. 2017; Katou et al. 2003; Byun et al. 2005). Helicase-polymerase uncoupling typically results in the activation of the intra-S-phase checkpoint (Byun et al. 2005). Helicase-polymerase uncoupling functions as a common consequence and signal to the intra-S-phase checkpoint across these mechanistically distinct perturbations to genome replication. Whether the uncoupled helicase also stalls through a common mechanism remains unclear. Rad53 appears to have a regulatory role in limiting DNA unwinding following uncoupling of the replisome in response to DNA damage or hydroxyurea (Lopes et al. 2006; Gan et al. 2017; Devbhandari and Remus 2020). Recent *in vitro* work in the context of UV lesions has suggested that the helicase slows and eventually stalls following uncoupling simply due to the absence of leading strand replication behind the helicase (Taylor and Yeeles 2019). In *E. coli,* this slowing of the helicase has also been observed when uncoupled from replication, and has been termed a “dead man’s switch” (Graham et al. 2017). The intrinsic or extrinsic mechanisms that ultimately halt CMG progression through chromatin following uncoupling remain unresolved in the context of *in vivo* chromosomal DNA replication.

In studies of helicase activation and helicase-polymerase uncoupling, the following open questions remain: i) How is local chromatin impacted by helicase activation? Is this similar at all origins, or does it vary based on the characteristics of individual origins? ii) What factors influence helicase movement and processivity when uncoupled from replication *in vivo*? iii) How does the cell respond to helicase-polymerase uncoupling outside of the contexts of DNA damage and replication stress? Can cells repair any unreplicated gaps left by this uncoupling? Using a conditional genetic system to target the function of polymerase α along with nucleotide-resolution genomic assays, we have captured helicase activation within chromatin at origins genome-wide, which has also allowed us to study the consequences of helicase-polymerase uncoupling *in vivo* in the absence of external damaging agents. Given the conservation of the CMG complex and its activity from the budding yeast to mammals (Aves et al. 2012), our results likely have implications for helicase activation and helicase-polymerase uncoupling within chromatin context in all eukaryotes.

## Results

### Capturing the active CMG complex at start sites of DNA replication

The formation of the CMG complex, helicase activation, DNA unwinding and initiation of DNA replication by polymerase α priming are tightly coupled events that must occur in the context of chromatin. To better understand the chromatin dynamics associated with the transition to an active CMG complex prior to initiation of DNA replication, we sought to block the priming of leading and lagging strand DNA replication by DNA polymerase α. In the absence of Pol α, the CMG complex still assembles at origins and can unwind DNA (Watase et al. 2012; Kanke et al. 2012; Van Deursen et al. 2012; Miyazawa-Onami et al. 2017; Douglas et al. 2018).

To prevent priming by Pol α and thus inhibit the initiation of DNA replication, we used temperature-sensitive alleles (*cdc17-1, cd17-2*) of the catalytic domain of Pol α (Hartwell et al. 1973; Lucchini et al. 1990) in combination with the Anchor Away system to abrogate Pol α function (Haruki et al. 2008). Upon rapamycin addition, the Anchor Away system rapidly depletes a protein of interest from the nucleus using components of the mTOR pathway. We initially characterized combinations of these alleles across restrictive conditions (temperature and/or rapamycin addition). (Figure 1A). Cells with the combination of both temperature-sensitive mutations and an Anchor Away tag, *cdc17-1,2-FRB*, exhibit the greatest sensitivity to the restrictive conditions (37°C + rapamycin).

**Figure 1.**
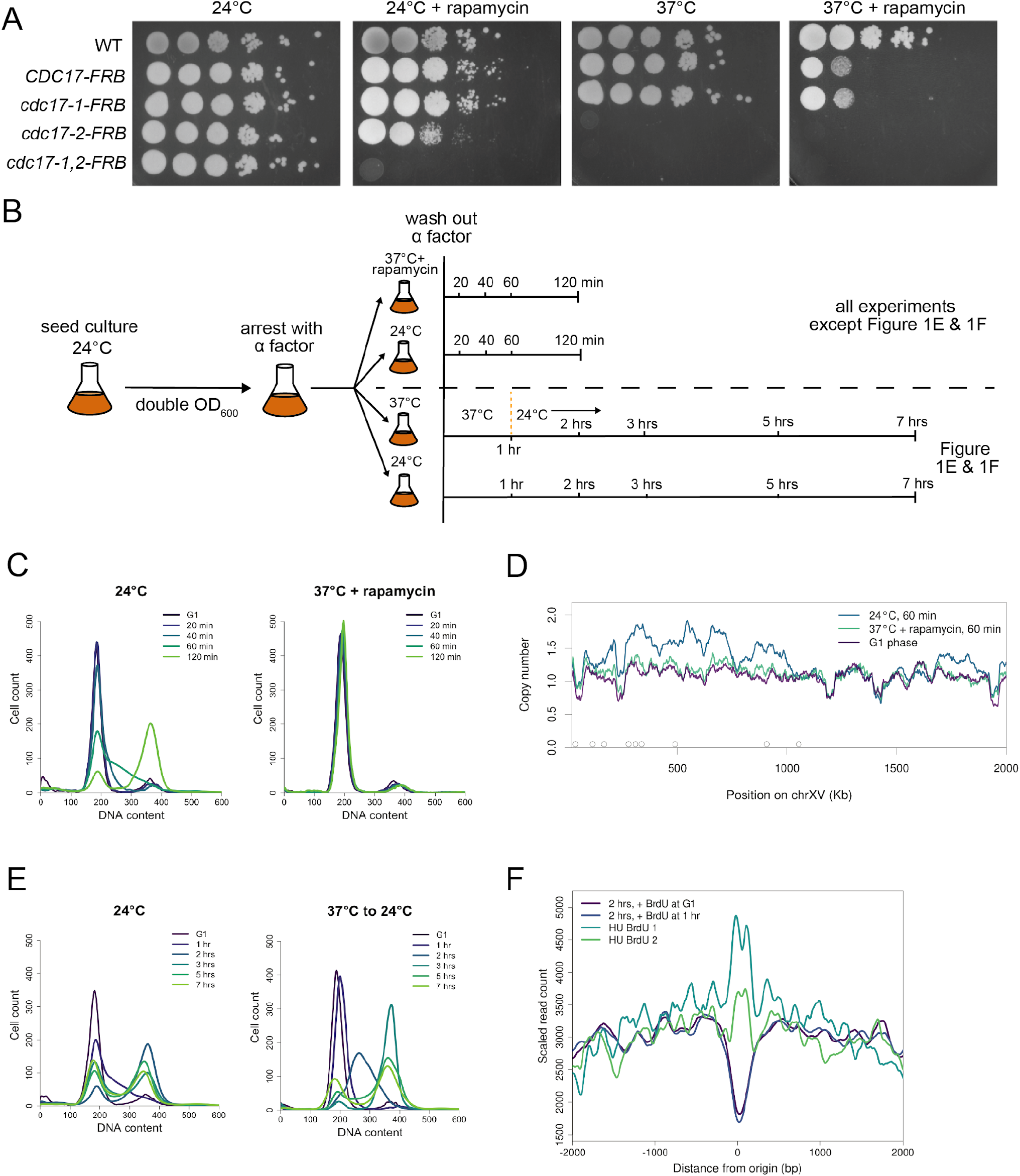
*cdc17-1,2*-*FRB* cells arrest prior to replication under restrictive conditions and recover from prolonged arrest with a delay in G2/M phase. (**A**) Spot assay across permissive and restrictive conditions for a series of *cdc17* alleles. (**B**) Schematic of experimental designs for arrest and recovery experiments. (**C**) DNA content measured by flow cytometry in *cdc17-1,2-FRB* cells under permissive (24°C) and restrictive conditions (37°C + rapamycin) during G1 phase and up to 120 minutes after release from G1. (**D**) Copy number across chromosome XV during G1 or 60 minutes after release from G1, under permissive (24°C) or restrictive (37°C + rapamycin) conditions. Open circles along the x-axis denote the positions of early origins. (**E**) DNA content measured by flow cytometry for *cdc17-1,2-FRB* cells under permissive (24°C) and restrictive conditions (37°C). Cells were synchronized in G1 phase and released for 7 hours, either with 1 hour at 37°C and 6 hrs at 24°C or 7 hrs at 24°C. (**F**) Aggregate BrdU enrichment at early origins. Samples were taken at the 1 hr 24°C recovery time point shown in **Figure 1B**, with BrdU added either upon release from G1 or upon return to 24°C. Hydroxyurea (HU) samples are WT cells released from G1 into HU for 2 hours at 30°C.

To ensure that DNA replication was completely inhibited by our approaches to remove Pol α function, we assessed the progression of *cdc17-1,2-FRB* cells through the cell cycle using flow cytometry. Cells were synchronized in G1 phase using α-factor, and monitored for 2 hours following release from G1 for entry into S-phase (Fig 1B & C). 20 minutes after release from G1 synchronization, cells still have G1 DNA content in both permissive and restrictive conditions, indicating that they are recovering from the α-factor arrest. At permissive conditions (24°C), cells begin to enter S-phase by 40 minutes. At 60 minutes after release from α-factor, most cells are in G2 phase, and by 2 hours they return to an asynchronous cycling pattern. Under restrictive conditions (37°C + rapamycin), cells remain arrested with DNA content equivalent to G1 phase even after 2 hours suggesting that no or minimal chromosomal replication occurs under restrictive conditions. To confirm this observation, we examined copy number genome-wide to determine whether any limited genome duplication was occurring (Figure 1D). Plotting copy number across Chromosome XV, we did not observe a copy number increase at early origins 60 minutes after cells were released from G1 arrest at 37°C with rapamycin. In contrast, at 24°C copy number increases approximately 50% at early origins 60 minutes after release from G1 phase. Taken in combination with our flow cytometry data, these experiments suggest that no or extremely minimal replication is occurring, making *cdc17-1,2-FRB* cells a useful *in vivo* model of helicase activation prior to the onset of replication.

In order to assess whether restoration of Pol α would allow recovery of replication following a prolonged arrest at 37°C, we synchronized cells as shown in Figure 1B and released them into 24°C or 37°C. Rapamycin was not used in the recovery experiments due to the difficulty in reversing the sequestration of the Anchor Away-tagged protein in the cytosol. After 1 hour, we returned cells to permissive conditions (24°C), and followed their progression through the cell cycle (Figure 1E). Control cells that were never exposed to the restrictive conditions progress through S-phase and G2/M phase following release from α-factor. In contrast, cells that experienced the restrictive temperature exhibit a prolonged S-phase and an arrest at G2/M phase; however, by 6 hours the cells eventually return to an asynchronous cell cycle distribution. Thus, loss of Pol α function causes a tight, but reversible arrest prior to replication. The subsequent G2 phase arrest experienced by cells recovering Pol α function suggests activation of the G2/M checkpoint to allow for completion of DNA replication and/or repair of DNA damage.

In the absence of DNA replication priming, the Mcm2-7 helicase may become active and start to unwind origin DNA, which has the potential to result in unreplicated gaps at replication origins. In order to assess whether there are any unreplicated gaps or disruptions to the replication program that might cause activation of the G2/M checkpoint following recovery, we labeled nascent DNA with the thymidine analog bromodeoxyuridine (BrdU) to assess the extent of DNA replication during the experiment (Figure 1F). BrdU incorporation into nascent DNA was detected by BrdU immunoprecipitation and sequencing (Viggiani and Aparicio 2006; Ryba et al. 2011). BrdU was added either upon release from G1 while cells were still at 37°C, or upon their return to 24°C (1 hr after G1). If there is helicase activation and unwinding in the absence of DNA replication, then when permissive conditions are restored, we would expect BrdU incorporation to be detected at sequences flanking or distal to replication origins, but depleted at the origin itself. Alternatively, if there is no helicase activation or unwinding, then we would expect uniform BrdU incorporation across origin sequences. We found a marked depletion for BrdU incorporation at replication origins, suggesting that delay in S-phase progression and G2/M arrest is due to the presence of unreplicated gaps.

It is formally possible that there was limited DNA replication at the non-permissive temperature which would result in the incorporation of thymidine nucleotides instead of BrdU at the origin proximal sequences. To exclude this possibility, we also added BrdU at the non-permissive temperature (37°C) following release from α-factor. The origin specific depletion of BrdU at the non-permissive and permissive temperature was identical (Figure 1F, purple and dark blue lines), consistent with the persistence of unreplicated gaps at replication origins. As an additional control, we also examined enrichment at early origins from BrdU-IP-seq experiments in which WT cells were released into HU for 2 hours. There is a local maximum in enrichment at early origins across both of these BrdU experiments, presumably due to favorable incorporation of the thymidine analog BrdU at AT-rich origin sequences. Overall, we conclude that in *cdc17-1,2-FRB* cells, there is likely no replication at restrictive conditions, and that an unreplicated gap is left at the origin when cells recover from the absence of Pol α, necessitating a delay in G2/M phase to repair these gaps. If the helicase can unwind DNA prior to the onset of replication, it could progress away from origins and leave behind the observed gap once replication begins. We thus sought to investigate the state of replication start sites under restrictive conditions in *cdc17-1,2-FRB* cells, beginning with chromatin structure.

### Origin chromatin is disrupted in the absence of Cdc17 function

Helicase activation has been modeled *in vitro* and at single origins *in vivo* (Miyazawa-Onami et al. 2017; Douglas et al. 2018). However, it has never been assessed in high resolution at origins across the genome. Additionally, despite the importance of local chromatin in origin function, the chromatin dynamics that accompany helicase activation have never been examined. We used a factor-agnostic micrococcal nuclease (MNase) chromatin profiling assay to study chromatin dynamics during helicase activation (Henikoff et al. 2011). To collect chromatin samples for this assay, we used the experimental design shown in Figure 1B, harvesting cells in G1 and at 20, 40, and 60 minutes after release from G1, both at 24°C and 37°C with rapamycin. Based on the FACS analysis (Figure 1C), cells are recovered from G1 arrest and in S phase by 40 minutes following release from α-factor. The resulting chromatin samples were digested with MNase, and DNA fragments between 20-250 bp in length were recovered. By plotting these fragment lengths as a function of chromosomal position, we can infer the positions of both nucleosomes and small DNA-bound proteins, such as replication and transcription factors. For example, during S-phase (both at 24°C and 37°C + rapamycin) we observe new small factors, likely transcriptional regulators, binding upstream of the gene *RNR1*, which encodes a ribonucleotide reductase subunit (Fig S1A). Additionally, chromatin structure is disrupted throughout the gene body, consistent with elevated transcription of *RNR1* in S-phase (Elledge and Davis 1990).

Based on the extensive DNA unwinding observed *in vitro* in studies of helicase-polymerase uncoupling and the transcriptional chromatin dynamics our assay can capture, we predicted that helicase activation and DNA unwinding would specifically disorganize chromatin structure around origins. We surveyed the genome for disorganized chromatin structure at origins under the restrictive conditions following release from α-factor, and identified specific origins that exhibited a time-dependent change in chromatin organization as they entered S-phase. For example, the early-activating origin ARS922 on chromosome IX exhibits precisely positioned nucleosomes flanking the origin and an ORC-dependent small DNA-binding factor footprint (Belsky et al. 2015) at the ACS in G1 and 20 minutes following release from α-factor (Figure 2A). However, by 40 minutes (when cells should be entering S-phase) we see a marked disorganization in the chromatin flanking the origin that persists for at least 1 hour following release. Specifically, small fragments <100 bp accumulate outside of known DNA-binding factor footprints (e.g. under nucleosome positions). Nucleosome-sized fragments are also detected between nucleosomes, and even intrude into the nucleosome-free region at the origin center. These changes may indicate loss of nucleosome positioning, additional small factors binding to DNA, and/or the presence of subnucleosomal structures (Rhee et al. 2014; Ramachandran et al. 2017). However, other origins, such as ARS405.5, a late origin on chromosome IV, have a static chromatin organization across all time points (Fig 2B). Additionally, control loci, such as binding sites for the transcription factor Abf1, also remain unchanged (Fig. S1B). We conclude that there is an origin-specific chromatin disorganization occuring in the absence of replication upon release of cells from G1 phase, and that this chromatin change is limited to a subset of replication start sites.

**Figure 2.**
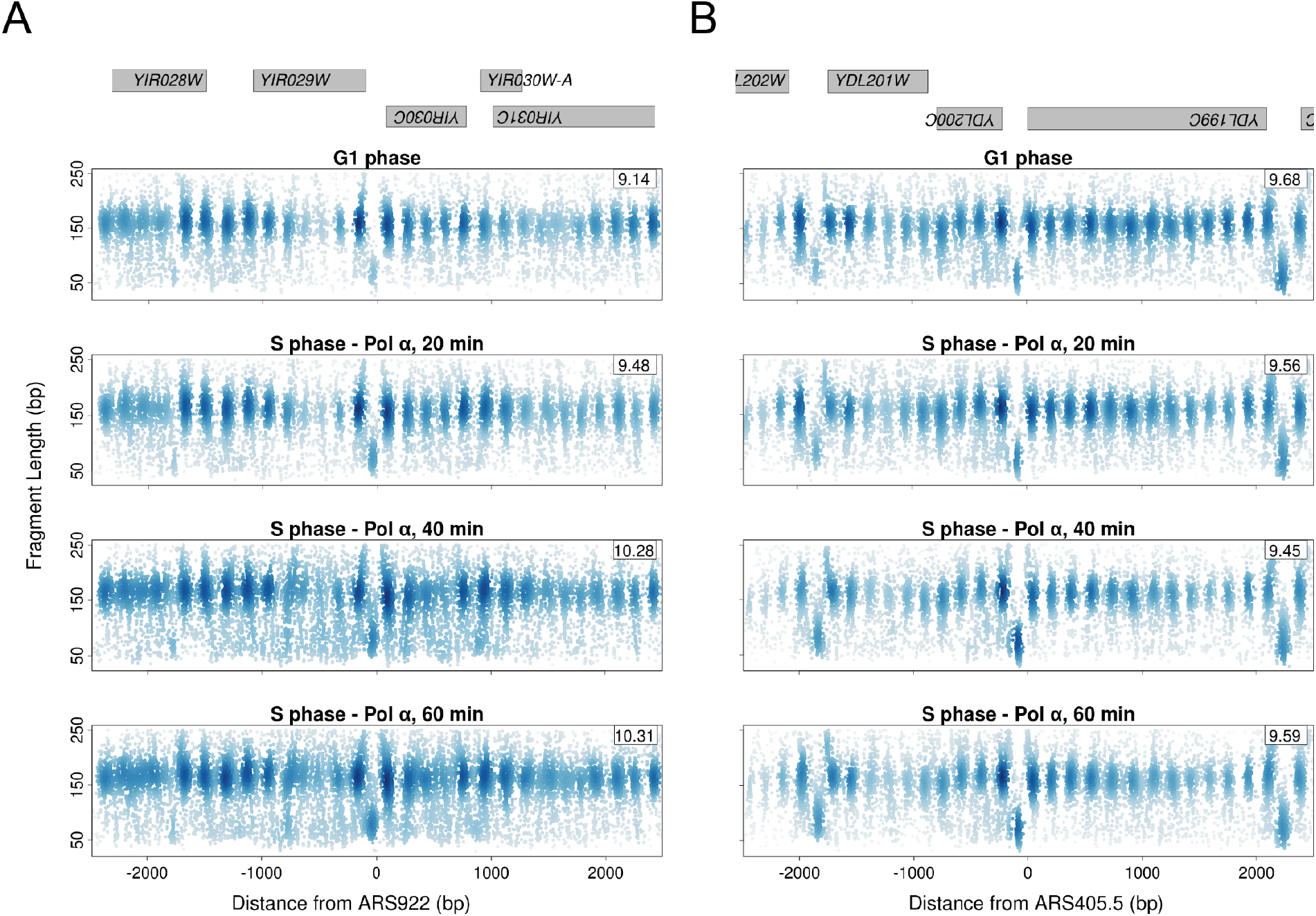
Chromatin disorganization occurs at select replication origins under restrictive conditions in *cdc17-1,2*-*FRB* cells. (**A**) Chromatin profiles for ARS922 in *cdc17-1,2*-*FRB* cells at 37°C + rapamycin, in G1 phase and 20, 40 and 60 minutes following release from G1. Fragment midpoints are plotted by fragment length and genomic position relative to the origin. Shading of each point is determined by a 2D kernel density estimate. Genes bodies are shown in gray across the top. The 2D Shannon entropy estimate in bits (log2) for the 1 Kb window around the origin is inset on each panel. (**B**) Chromatin profiles for ARS405.5 in *cdc17-1,2*-*FRB* cells at 37°C + rapamycin, in G1 phase and up to 60 minutes after release from G1.

We hypothesized that if the observed chromatin disorganization resulted from the presence of active CMG complexes at origins, it should be limited to early, efficient origins. In the absence of replication, limiting initiation factors necessary to form CMGs would remain trapped at these sites, unable to activate other Mcm2-7 double hexamers (Mantiero et al. 2011; Tanaka et al. 2011). In this model, late origins will not experience chromatin changes, either solely due to the absence of limiting pre-initiation complex factors or due to the additional suppression of late origins by the intra-S-phase checkpoint. To agnostically identify disorganized origins, we developed an entropy score based on information theory to holistically capture chromatin organization (Tran et al. 2020). In organized chromatin, fragment midpoints resulting from our MNase assay tend to be well-organized at small factor (e.g. ORC) binding sites or nucleosome dyads, while midpoint fragments recovered from disorganized chromatin are more randomly distributed. We estimated the 2D Shannon entropy around origins based on the distribution of fragment midpoints surrounding each origin, and used the difference in entropy between G1 and 40 minute chromatin as a discriminating score (see Suppl. Methods). A subset of 239 replication start sites that bind ORC during G1 & G2 phase was used in this analysis (Belsky et al. 2015). We assigned each origin an entropy score and plotted the distribution of these scores (Fig. 3A). Origins were classified as early activating if they activated in the presence of hydroxyurea and all others were designated as late origins (Belsky et al. 2015). The bimodal distribution of origin entropy scores demonstrates that early-firing origins have a greater entropy score, and thus more chromatin disorganization, than origins that activate later in S-phase.

**Figure 3.**
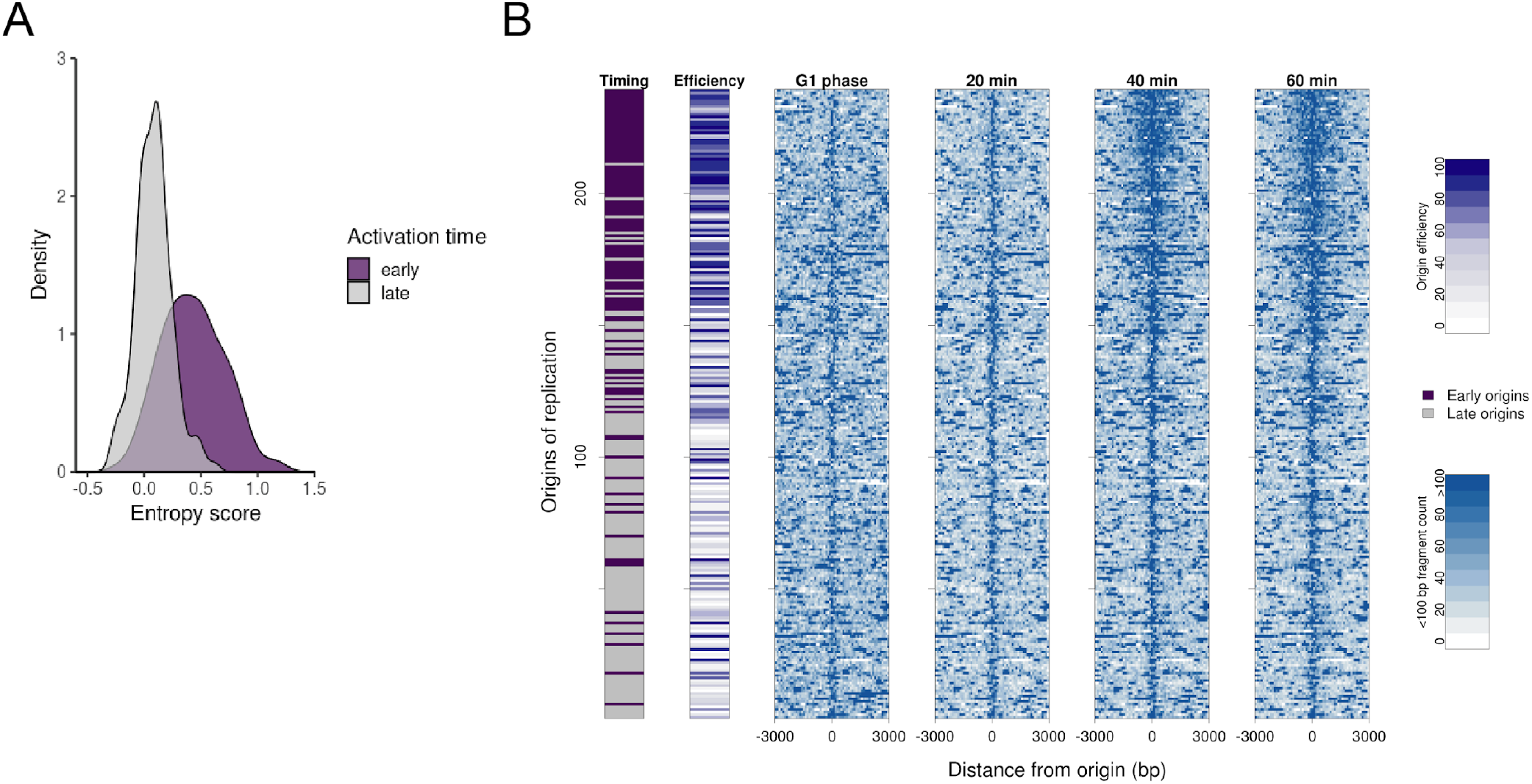
Early, efficient origins have disorganized chromatin extending up to 1 Kb away from the origin under restrictive conditions. (**A**) Distribution of entropy scores across early and late origins. The score represents the entropy difference between G1 phase chromatin and chromatin 40 minutes after release from G1 under restrictive conditions. (**B**) Heatmap of 239 replication origins in *cdc17-1,2-FRB* cells at 37°C + rapamycin, ordered by decreasing entropy score. The midpoints of <100 bp fragments resulting from MNase digestion were counted around each origin, and are plotted in 100 bp bins. The activation time and efficiency for each origin is annotated on the left. Origin timing indicates a binary early or late designation based on whether or not an origin activates in HU. Time labels reflect minutes following release from G1 phase.

We next ordered origins based on this score and visualized the increased entropy by only plotting those fragments with a length of <100 bp which were overrepresented in the disorganized chromatin (Fig 3B). Timing and efficiency annotations for each origin further confirm that early-activating, efficient origins have disorganized chromatin relative to G1 phase (McGuffee et al. 2013; Belsky et al. 2015). In contrast, late, inefficient origins retain their G1 chromatin structure. The increase in chromatin disorganization as reflected by the entropy score extends up to 1 Kb on either side of origins. As observed at ARS922 (Figure 2A), chromatin disorganization does not occur until 40 minutes after release from G1 arrest, and it persists at 60 minutes. The increased entropy observed at early, efficient origins only occurs under restrictive conditions at origins; it is not present at origins under permissive conditions (Figure S2A) or at Abf1 binding sites (Figure S2B). These results support that the chromatin disorganization we observe is dependent on the formation of the pre-initiation complex, since the phenomena is limited to early, efficient origins.

### CMG movement and ssDNA accumulation define the extent of chromatin disorganization around origins

The observation that chromatin disorganization is limited to early, efficient origins in a Pol α conditional mutant suggests that helicase activation and DNA unwinding might be responsible for these chromatin dynamics. If this is the case, Mcm2-7 would be detected outside the origin center where it is loaded in G1 phase, likely 1 Kb from the origin center. To test this hypothesis, we performed Mcm2-7 ChIP-seq across the same time course in *cdc17-1,2-FRB* cells and examined Mcm2-7 ChIP enrichment at sequences surrounding origins of replication. To visualize the data, we ordered origins by their chromatin disorganization score as in Figure 3B. In G1 phase, Mcm2-7 is located at the origin center (ACS) across all origins (Fig 4A). Beginning 40 minutes after release from G1 synchronization, Mcm2-7 is enriched approximately 1 Kb up and downstream from the same subset of early, efficient origins at which we observe chromatin disorganization (Figure 4B). This pattern of Mcm2-7 enrichment only occurs under restrictive conditions (Fig S3A & B). The bi-directional enrichment of Mcm2-7 suggests the double hexamer has separated, which occurs when the CMG complexes are fully formed (Douglas et al. 2018). To confirm that the complete CMG complex, rather than Mcm2-7 alone, is moving away from the origin, we performed ChIP-seq for FLAG-tagged Pol2, a subunit of polymerase ε (Fig. 4C). Pol ε is recruited to origins during pre-initiation complex formation following S-CDK phosphorylation along with the GINS complex, and travels with the active CMG (Heller et al. 2011; Sengupta et al. 2013; Yu et al. 2014). We observe Pol ε enrichment in the same pattern as the Mcm2-7 complex flanking the origin 1 Kb up and downstream, suggesting the intact CMG complex is moving bidirectionally away from these origins in the absence of Pol α. These results support our hypothesis that the active holo-helicase complex is able to unwind DNA and perturb chromatin structure prior to the onset of replication.

**Figure 4.**
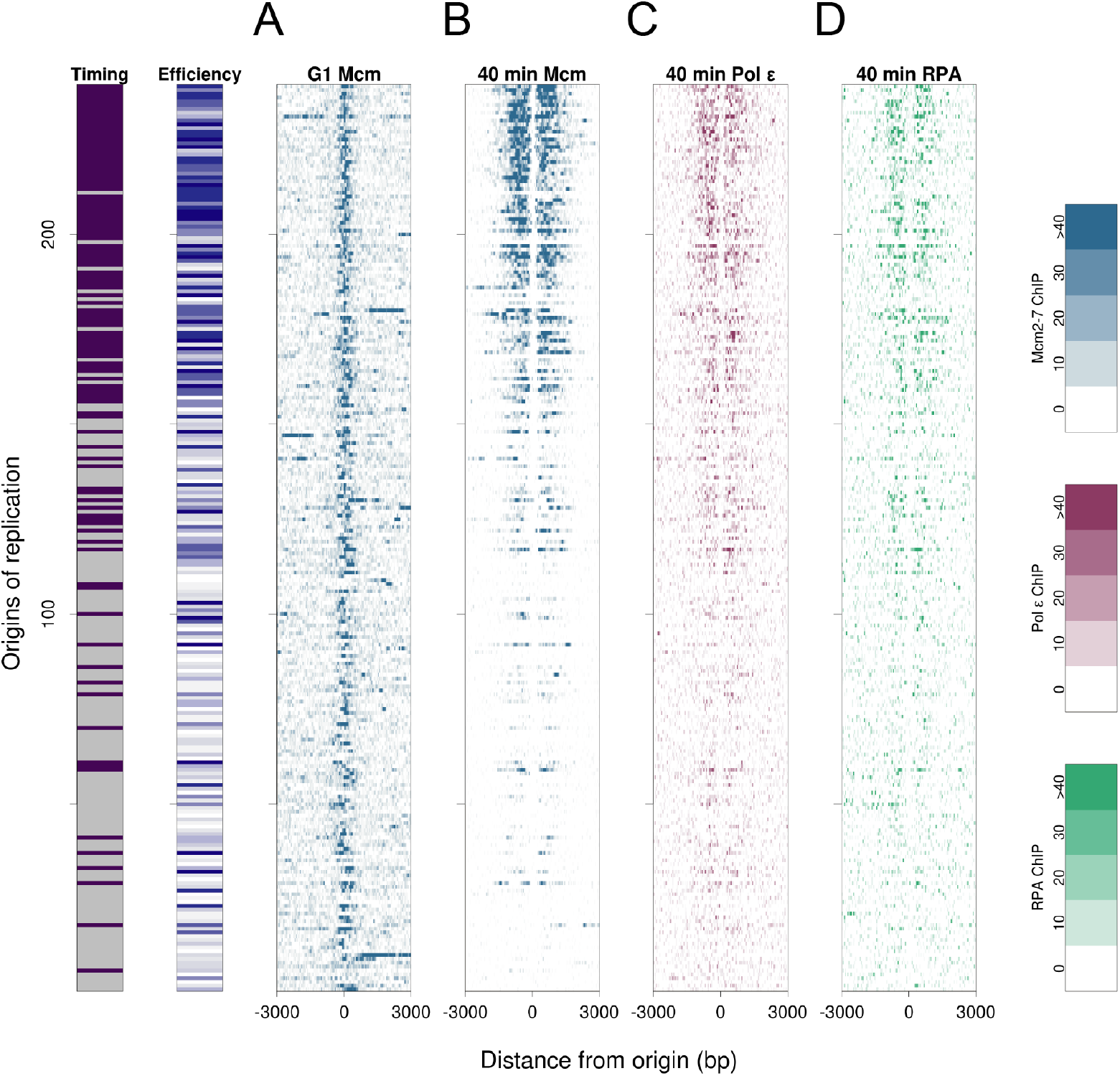
CMG movement and ssDNA accumulation define the extent of chromatin disorganization. (**A**) Mcm2-7 ChIP-seq enrichment across origins during G1 phase and (**B**) 40 min after G1 at 37°C + rapamycin in *cdc17-1,2-FRB* cells, ordered by entropy score as in **Fig. 3**. G1 enrichment has been subtracted from the 40 min enrichment shown. Enrichment plotted in 50 bp bins. (**C**) Pol2 (Pol E subunit) ChIP-seq enrichment at origins 40 minutes after G1 phase, normalized to G1 Pol2 enrichment as in **A**. (**D**) Rfa1 (RPA subunit) ChIP-seq enrichment at origins 40 minutes after G1, normalized to G1 Rfa1 enrichment as in **A**.

During G1 the Mcm2-7 double hexamer can translocate along double-stranded DNA (dsDNA) when pushed by an RNA polymerase (Gros et al. 2015). Thus, it is possible that the CMG complex could be moved in a similar way after double hexamer separation without unwinding DNA, although this has not been previously observed in the context of the CMG complex. To determine if ssDNA was produced by CMG movement, we performed ChIP-seq using a tagged allele of *RFA1*, a subunit of the ssDNA binding complex RPA. Plotting Rfa1 enrichment at 40 minutes alongside the Mcm2-7 and Pol ε ChIP data, we observe that Rfa1 is bound to DNA in a similar pattern flanking the same set of origins (Figure 4D). The enrichment of Rfa1 at sequences flanking early origins is specific to the restrictive conditions and does not occur under permissive conditions (Figure S3C). Our data suggests active CMG complexes are unwinding DNA at a subset of origins, and this ssDNA persists long enough for us to observe RPA binding. This unwinding appears to be limited to a 1 Kb window around origins, suggesting that there are mechanisms that halt CMG movement when uncoupled from replication.

### The intra-S-phase checkpoint does not regulate CMG progression in the absence of replication

When tracts of ssDNA accumulate following replication stress, RPA binds to this ssDNA. At a certain threshold, this ssDNA-RPA signal activates the DNA replication arm of the intra-S-phase checkpoint (Pardo et al. 2017). The DNA replication checkpoint is mediated through Mrc1, and leads to the phosphorylation and activation of the effector kinase Rad53. Rad53 suppresses late origin firing, along with other actions that halt further replication, and has recently been shown to regulate the replisome during helicase-polymerase uncoupling *in vitro* (Tercero and Diffley 2001; Devbhandari and Remus 2020).

The persistent RPA accumulation we observe in *cdc17-1,2-FRB* cells, which occurs at early and not late origins, suggests activation of the intra-S-phase checkpoint. We hypothesized that Mcm2-7 movement might be regulated by the activated intra-S-phase checkpoint in the absence of replication. In the absence of the checkpoint, we predicted that Mcm2-7 might continue to progress further than 1 Kb from origins, and that Mcm2-7 movement would occur at both late and early origins. To test these predictions, we constructed *cdc17-1,2-FRB MRC1-FRB* cells to examine helicase movement in the absence of the intra-S-phase checkpoint; similar to *cdc17-1,2-FRB* cells, these cells arrest with G1 DNA content under restrictive conditions (Fig. S4A & S4B). To assess checkpoint status in *cdc17-1,2-FRB* cells alone and in combination with *MRC1-FRB*, we performed immunoblots for Rad53 in both strains (Fig. 5A). In *cdc17-1,2-FRB* cells, Rad53 is phosphorylated beginning 40 minutes after release from G1 arrest, corresponding to the time we observe chromatin disorganization, CMG movement, and RPA binding around origins. Thus, the amount of ssDNA observed in Figure 4 is sufficient to activate the intra-S-phase checkpoint. Rad53 phosphorylation does not occur at any time point in *cdc17-1,2-FRB MRC1-FRB* cells under restrictive conditions, indicating intra-S-phase checkpoint activation is blocked by depletion of Mrc1 from the nucleus. We performed our MNase chromatin profiling assay and Mcm2-7 ChIP-seq in *cdc17-1,2-FRB MRC1-FRB* cells to assess Mcm2-7 movement and chromatin disorganization in the absence of the checkpoint. To compare Mcm2-7 ChIP enrichment across origins, we ordered origins based on the Mcm2-7 ChIP signal in *cdc17-1,2-FRB* cells. As expected, Mcm2-7 occupancy and movement in the absence of the intra-S-phase checkpoint is distributed across a larger set of origins compared to cells with the checkpoint intact (Fig. 5B). However, Mcm2-7 movement is still limited to 1 Kb on either side of the origins.

**Figure 5.**
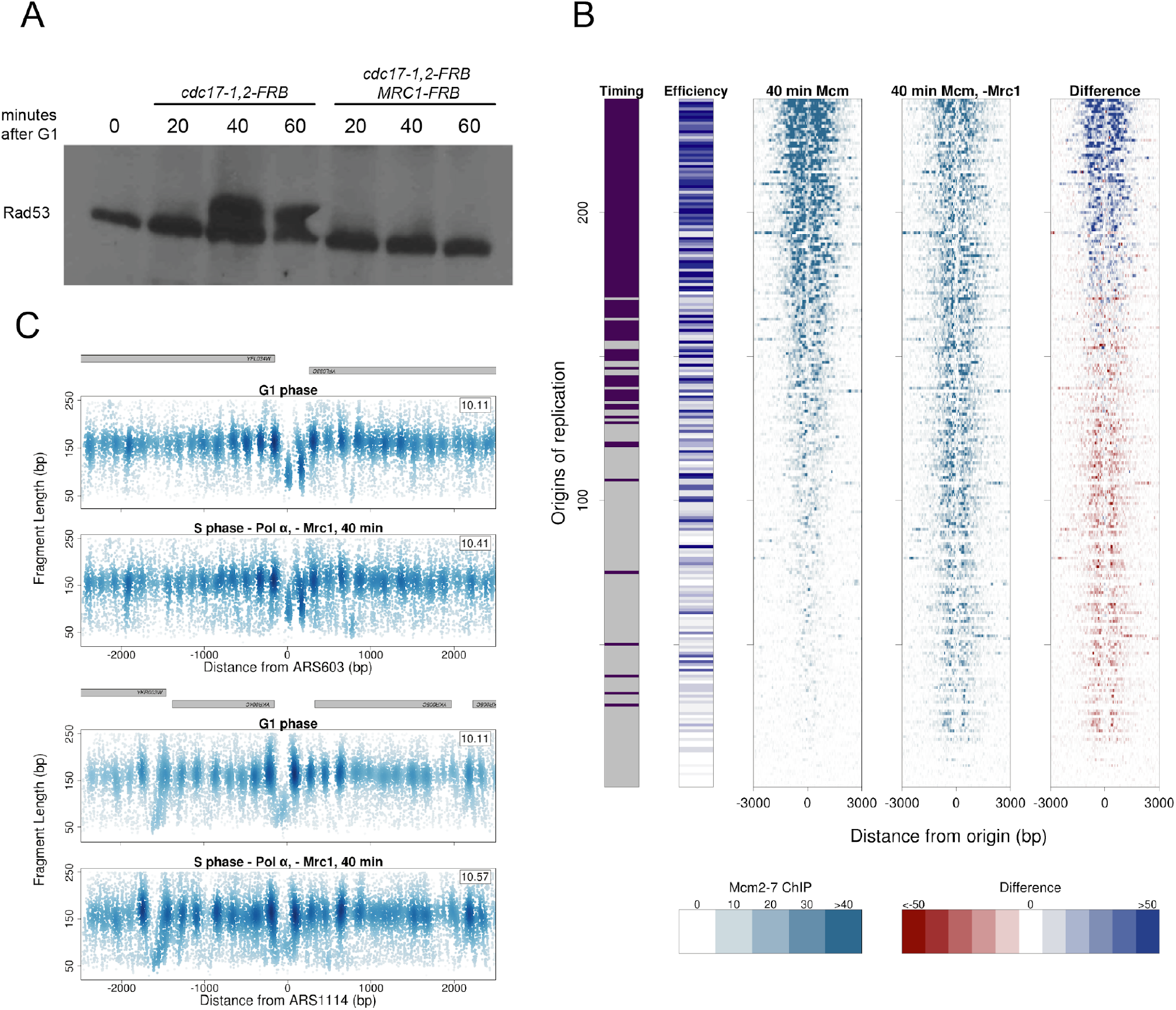
The intra-S-phase checkpoint is activated in the absence of Cdc17 function, but does not influence the extent of Mcm2-7 movement. (**A**) Western blot for Rad53 in *cdc17-1,2-FRB* and *cdc17-1,2-FRB MRC1-FRB* cells, at 37°C + rapamycin following release from G1 synchronization. The upper band indicates phosphorylated Rad53. (**B**) Heatmap of Mcm2-7 ChIP-seq enrichment in 50 bp bins at origins in *cdc17-1,2*-*FRB* cells at 37°C + rapamycin, with and without *MRC1-FRB*. Origins are ordered by decreasing Mcm2-7 enrichment level at 40 min in *cdc17-1,2-FRB* cells. The difference panel was calculated by subtracting the Mcm2-7 signal in *cdc17-1,2*-*FRB MRC1-FRB* from the Mcm2-7 signal in *cdc17-1,2*-*FRB.* (**C**) Chromatin profiles for early origin ARS603 and late origin ARS1114 in *cdc17-1,2*-FRB *MRC1-FRB* cells, in G1 phase and 40 minutes after release from G1 at 37°C + rapamycin. The entropy value for +/−1 Kb around each origin is shown in the top left of each panel. Gene bodies are depicted in gray at the top of each plot. Points are shaded as in **Figure 2**.

When we analyzed origin chromatin in our MNase data using the entropy score, we found that chromatin disorganization, likely due to the presence of an active helicase, now occurs at both early and late replicating origins. (Fig. S4C). For example, ARS603 is a late origin that gained disorganized chromatin 1 Kb in *cdc17-1,2-FRB MRC1-FRB* cells (Fig. 5C). Early origins, such as ARS1114, still exhibit chromatin disorganization. Thus, the intra-S-phase checkpoint is involved in halting the activation of late replication origins in the absence of replication, similar to its role under replication stress conditions. Though the helicase is active at most origins, the same limits on Mcm2-7 unwinding and movement persist suggesting that other factors must influence helicase movement and DNA unwinding in the absence of replication.

### CMG movement stalls in a sequence-dependent manner

Since the intra-S-phase checkpoint is activated by Mcm2-7-dependent DNA unwinding in the absence of replication, but does not appear to limit the movement of Mcm2-7, we speculated that local chromatin features or sequence elements surrounding origins may hinder helicase progression. We first examined the local chromatin environment surrounding origins. If nucleosomes act as barrier elements, Mcm2-7 peaks should accumulate at these features. We selected highly enriched Mcm2-7 peaks on either side of the origins at 40 minutes. However, when we looked at nucleosome and Mcm2-7 occupancy, we did not observe specific enrichment of Mcm2-7 at nucleosome dyads or in linker regions (Fig. S5). Thus, nucleosome positioning does not appear to be the ultimate reason Mcm2-7 stalls, though it may still have some inhibitory effect on DNA unwinding.

We next examined the sequence content surrounding Mcm2-7 peaks. Replication origins in *S. cerevisiae* share an AT-rich DNA unwinding element (Umek and Kowalski 1988), and therefore GC content increases with distance from the origin until it reaches the genome-wide average. If sequence context inhibits origin unwinding, Mcm2-7 peaks would likely stall outside the AT-rich region, which is easier to unwind due to its lower thermal stability (Umek and Kowalski 1988; Natale et al. 1992; Ak and Benham 2005). We first plotted aggregate GC content and Mcm2-7 ChIP enrichment surrounding origins in *cdc17-1,2-FRB* cells (Fig 6A). GC content at replication origins decreases from the genome-wide average of 37% to 20%. Mcm2-7 is strongly enriched just beyond the edges of this AT-rich origin region where the GC content returns to the genome-wide average. We also observe that the Mcm2-7 enrichment pattern closely aligns with the distribution of nascent BrdU-labeled DNA following recovery from 37°C, first shown in Figure 1F. Together, these results suggest that replication begins where Mcm2-7 stalls leaving behind an unreplicated gap.

**Figure 6.**
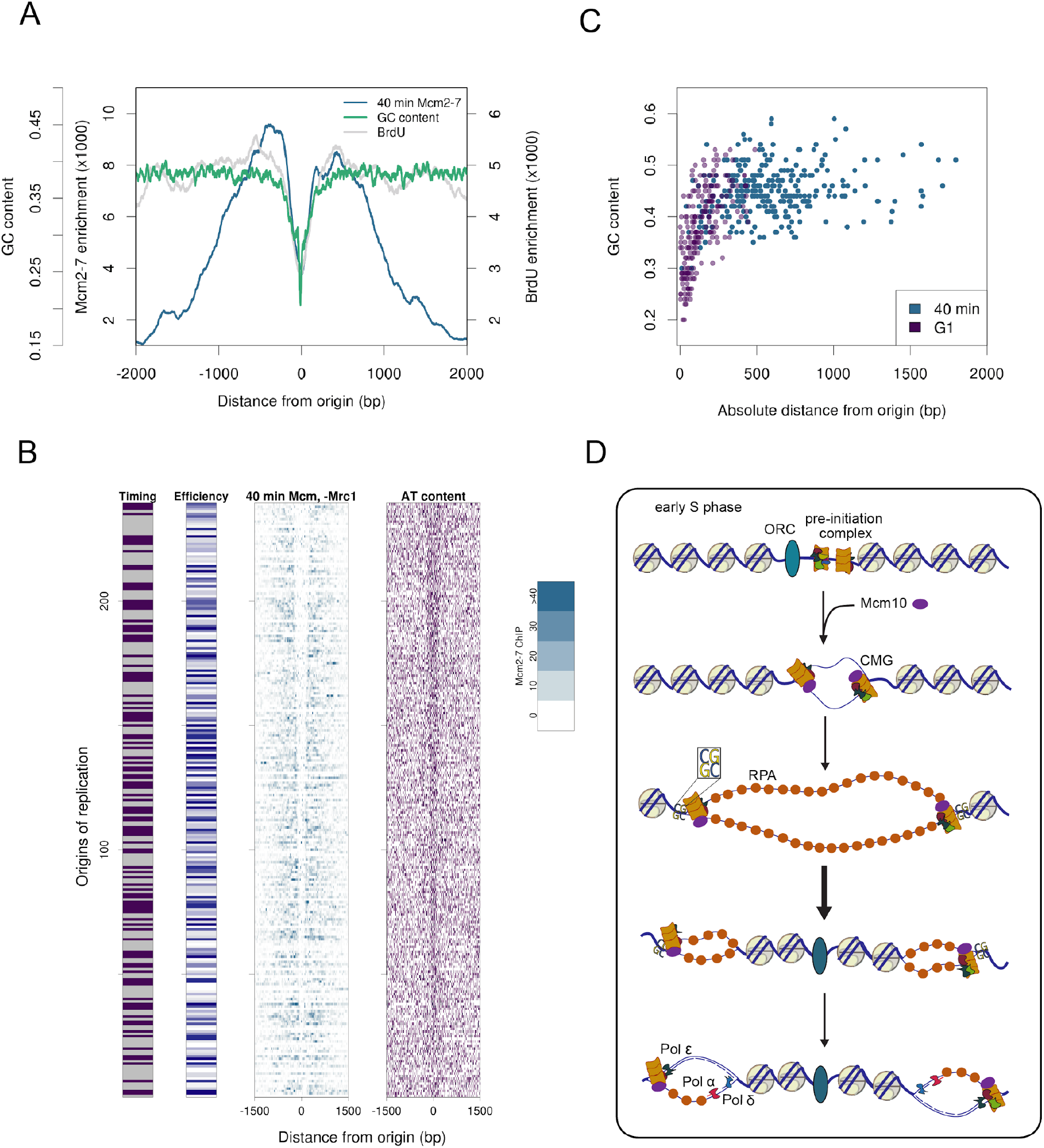
The active CMG complex unwinds the AT-rich origin region and stalls as it moves through sequences with higher GC content. (**A**) Aggregate GC content and Mcm2-7 ChIP enrichment at 37°C + rapamycin at 239 origins in *cdc17-1,2*-*FRB* cells. Aggregate BrdU-IP-seq signal (not scaled) from **Figure 1F**, 2 hrs after BrdU addition and release from G1, is shown for comparison. GC content calculated in 30 bp bins. (**B**) (*left*) Heatmap of Mcm2-7 ChIP-seq enrichment at origins 40 minutes after G1 under restrictive conditions in *cdc17-1,2*-*FRB MRC1-FRB* cells. Origins are ordered by decreasing length of the AT-rich region at origins. (*right*) AT content around origins. 3-mers containing only A/T are shaded purple; all other 3-mers are white. (**C**) GC content and distance from origin for each Mcm2-7 ChIP-seq peak in *cdc17-1,2*-*FRB* cells during G1 phase and after 40 minutes released from G1, 37°C + rapamycin. GC content is calculated from a 100 bp window around each Mcm2-7 peak. (**D**) Proposed model for helicase activation without replication and recovery following restoration of Pol α.

To further examine Mcm2-7 positioning relative to the origin sequence, we ordered origins based on the length of their AT-rich regions and plotted Mcm2-7 ChIP signal in *cdc17-1,2-FRB MRC1-FRB* cells (Fig 6B). Mcm2-7 is seemingly excluded from the AT-rich region, and accumulates abruptly when GC content increases. Together, our data indicate that active CMG complexes stall when encountering an increase in GC content when DNA unwinding is uncoupled from replication.

To more precisely compare Mcm2-7 movement and GC content, we plotted each Mcm2-7 peak by GC content and distance from the origin (Fig 6C). G1 phase Mcm2-7 peaks tend to be close to the origin, with a median GC content of 38%. Conversely, 40 minutes after release from G1 phase under restrictive conditions, Mcm2-7 peaks have a significantly higher median GC content of 45% (p-value < 2.2e-16, t-test), and are located farther from the origin center than the G1 peaks. Overall, we conclude that the active CMG moves away from the origin to regions of higher GC content after release from G1 arrest in the absence of replication.

## Discussion

Replication start site selection, licensing, and activation occur within the local chromatin environment, and we have previously addressed how ORC binding and Mcm2-7 loading impact origin chromatin (Eaton et al. 2010; Belsky et al. 2015). Mechanistic studies of helicase activation have established that the two CMG complexes move apart at the origin prior to replication initiation (Douglas et al. 2018). The active helicase is also able to unwind kilobases of DNA when uncoupled from replication elongation (Walter and Newport 2000; Pacek and Walter 2004). Taken together, this evidence suggests that once active, the CMG has the potential to extensively unwind DNA and disrupt chromatin at the origin even in the absence of nascent strand synthesis. We have used multiple genomic approaches in a polymerase α conditional mutant to study both helicase activation and helicase-polymerase uncoupling within origin chromatin.

Activation of CMGs in the absence of replication results in chromatin disorganization at early, efficient origins characterized by loss of nucleosome positioning and an increase in entropy for fragments <100 base pairs. The increased entropy observed at replication origins in the absence of polymerase α function is analogous to the chromatin changes observed at highly transcribed genes upon induction, such as *RNR1*. The elevated entropy within gene bodies is driven by the continual unwinding by the RNA polymerase that disrupts nucleosome structure (Tran et al. 2020). In contrast, the increased chromatin disorganization and entropy at replication origins is likely driven by CMG-dependent DNA unwinding. Given the analogous chromatin dynamics at transcribed genes, it is likely that the observed dynamics are caused by changes in a common element, the nucleosome. At both sets of genomic loci, the small fragments likely reflect partially unwrapped nucleosomes or small structures such as tetrasomes, as has been postulated previously for fragment sizes less than 150 bp (Rhee et al. 2014; Ramachandran et al. 2017). Partially unwrapped nucleosomes would be more vulnerable to MNase digestion closer to the nucleosome dyad, resulting in shorter protected fragments, while subnucleosomal structures would simply protect fewer basepairs of DNA.

Despite the increase in entropy, well-positioned nucleosomes can still be discerned around the origin suggesting a mixed population of chromatin organizational states. In some cells, the helicase may have unwound less duplex DNA, leaving some nucleosomes well-positioned. In others, the helicase may not have been activated by pre-initiation complex factors, given the stochastic nature of the replication program. It is also energetically favorable that unwound, complementary single strands would eventually reanneal, allowing nucleosomes to be reset behind the helicase (Figure 6D). All of these possible states of unwinding and resetting will be captured by our chromatin profiling assay.

Chromatin disorganization, Mcm2-7 enrichment, and RPA enrichment at replication origins all support that the active CMG can uncouple from DNA replication and unwind duplex DNA up to 1 Kb from origins. Our prior work demonstrated that Mcm2-7 is loaded in the nucleosome-free region adjacent to either the upstream or downstream flanking nucleosome during G1 phase (Belsky et al. 2015). Interestingly, this asymmetric loading of the helicase near either the upstream or downstream nucleosome in G1 does not impact initial unwinding, which is symmetric with respect to the origin center. This suggests that the origin recognition complex does not pose a significant obstacle to the CMG, which can disrupt up to 6 nucleosomes on either side of the nucleosome-free region at the origin. Previous estimates of CMG movement in the absence of replication *in vivo* in budding yeast range widely from 100 bp up to 3 Kb, though 400-500 base pairs is the most common maximum estimate (Walter and Newport 2000; Sogo et al. 2002; Katou et al. 2003; Nedelcheva et al. 2005; Zellweger et al. 2015; Gan et al. 2017). This range of distances may be due to the different initial causes for helicase-polymerase uncoupling, which include dNTP depletion, multiple types of DNA lesions, and polymerase inhibition.

Mechanistic experiments *in vitro* support the model that after activation, CMG complexes establish replication forks traveling with the Mcm2-7 N-terminal domain first (Georgescu et al. 2017; Douglas et al. 2018). However, the Mcm2-7 double hexamer forms in G1 phase through the association of each Mcm2-7 complexes’ N-terminal domain, and thus this directionality requires the two CMG complexes to translocate past each other on ssDNA prior to initiation of replication. A consequence of this head-to-head model is more extensive DNA unwinding prior to replication initiation than if Mcm2-7 traveled with the opposite orientation at forks. Thus, limited helicase uncoupling at the origin may be a necessary feature for replication initiation.

Helicase-polymerase uncoupling in the absence of replication leaves behind unreplicated regions at origins when replication finally begins. Given that the location at which the Mcm2-7 complex stalls coincides with the location of BrdU incorporation following recovery, we hypothesize that in *cdc17-1,2*-*FRB* cells replication restarts outside the origin region (model in Figure 6D). Cells with limited levels of polymerase α experience increased genomic instability (Song et al. 2014), potentially due to ssDNA tracts generated by uncoupling of the leading and lagging strands. However, the unreplicated gaps we observe suggest that helicase-polymerase uncoupling at replication origins might contribute to this heightened instability as well. Prior to recovery, DNA unwound by the stalled CMG would likely reanneal, potentially aided by proteins that serve this role in recombination or repair, such as Rad52 (Shinohara et al. 1998). When polymerase α function is restored the CMG would resume unwinding DNA from where it stalled, and replication of the majority of the genome would begin. The resulting structure, two replicated double helices connected by an unreplicated section of reannealed parental strands at the origin, does not resemble any structure known to occur during homologous recombination or DNA repair pathways. There must exist a mechanism to repair this structure, given that recovering cells pause in G2/M phase and eventually re-enter the cell cycle. These gaps must first be recognized by a repair or checkpoint factor, and we expect that a helicase will be recruited to unwind the unreplicated strands. There are many known helicases that could serve this role, including Sgs1 and Srs2 (Crickard and Greene 2019). Once unwound, polymerase α could establish primers extended by Pol δ, and the nascent strands could ultimately be ligated together using mechanisms of Okazaki fragment maturation (Bell and Labib 2016).

Together our flow cytometry, copy number, and BrdU incorporation experiments support that *cdc17-1,2*-*FRB* cells do not initiate replication under restrictive conditions, but we still detect activation of the intra-S-phase checkpoint through Rad53 phosphorylation. The lack of Rad53 phosphorylation in *cdc17-1,2-FRB MRC1-FRB* suggests that intra-S-phase checkpoint activation occurs through the DNA replication arm of the checkpoint in this context, as opposed to the DNA repair arm mediated through Rad9 (Pardo et al. 2017). Canonically, the presence of ssDNA recruits the sensor kinase Mec1 (ATR in mammals), but it is not sufficient for Mec1 activation (Zou and Elledge 2003). The 9-1-1 complex, which recognizes ssDNA-dsDNA junctions, is one of the additional factors that contributes to activation of Mec1 (Majka et al. 2006; Majka and Burgers 2003). At minimum, a single primer is thought to be required for checkpoint activation, with additional primers amplifying activation (Byun et al. 2005; MacDougall et al. 2007; Van et al. 2010). However, if priming is absent in our system, there may be more flexibility in requirements for checkpoint activation than previously thought. Given that studies directly addressing the requirement of priming in checkpoint activation were done in *Xenopus*, this flexibility in checkpoint activation may be specific to budding yeast. In *S. cerevisiae*, additional factors such as Pol ε, play a role in the checkpoint, and the 9-1-1 complex may not be fully necessary for checkpoint function (Crabbé et al. 2010; García-Rodríguez et al. 2015).

Helicase-polymerase uncoupling has been described in various checkpoint mutants when exposed to DNA damage or hydroxyurea, and the effector kinase of the intra-S-phase checkpoint, Rad53, has been proposed as a key regulator of CMG progression in the absence of replication (Gan et al. 2017; Devbhandari and Remus 2020). In our experiments, Rad53 is phosphorylated in the absence of polymerase α function in *cdc17-1,2*-*FRB* cells, which confirms studies in other eukaryotes showing that the intra-S-phase checkpoint is activated in the presence of aphidicolin or polymerase α inhibition (Byun et al. 2005; Ercilla et al. 2020). We found that the intra-S-phase checkpoint does not ultimately regulate movement of the CMG complex when polymerase α function is perturbed. However, the context in which helicase-polymerase uncoupling occurs may modulate the role of Rad53 in regulation. For example, eSPAN experiments in *rad53-1* cells demonstrate that leading strand uncoupling in hydroxyurea can be almost entirely rescued by deletion of *SML1*, which inhibits ribonucleotide reductase RNR (Gan et al. 2017). Additionally, the same study found that Pol ε is more sensitive than Pol δ to dNTP levels. These results suggest that the role of Rad53 in regulation of dNTP pools is specific to uncoupling in cells exposed to hydroxyurea. *In vitro* reconstitution experiments have shown that an activated Rad53 can reduce the amount of DNA unwound by the CMG in the presence of aphidicolin, both at stalled forks and in the absence of replication (Devbhandari and Remus 2020). Given these results, Rad53 may be recruited to origins by the activation of the checkpoint in our experiments as well; however, Rad53 phosphorylation is not required for the helicase uncoupling we observe at DNA replication origins. Together these results suggest that there may be other extrinsic mechanisms that can stall the helicase.

Budding yeast origins, like some prokaryotic origins, have been shown to contain a broad AT-rich DNA unwinding element in addition to the shorter T-rich motif (Umek and Kowalski 1988; Kowalski and Eddy 1989). This DNA unwinding element is energetically easier to unwind into single strands, and likely serves to facilitate replication initiation (Natale et al. 1992; Ak and Benham 2005). As sequence content returns to the genome-wide average outside the origin region and becomes more difficult to denature, Mcm2-7 enrichment increases in *cdc17-1,2*-*FRB* under restrictive conditions, indicating the CMG is less processive and frequently stalls at sequences with elevated GC content. Though this pattern does not necessarily imply causation, previous studies support the model that replicative helicases are less processive following uncoupling with replication. Studies of the *E. coli* helicase and polymerases have established that when there are stochastic pauses in replication or if the helicase and polymerase are physically separated, the rate of helicase progression slows to prevent excessive uncoupling (Kim et al. 1996; Yeeles and Marians 2013; Graham et al. 2017). This phenomena has been named a biological “dead man’s switch” (Graham et al. 2017). It has recently been proposed that a similar mechanism exists in the eukaryotic replisome (Taylor and Yeeles 2019). When Mcm2-7 encounters a DNA lesion on the leading strand, the replisome slows down or stalls regardless of the type of lesion or composition of the replisome, suggesting there is an intrinsic mechanism that stalls the helicase (Sparks et al. 2019; Taylor and Yeeles 2019). Given that the CMG can bypass leading strand DNA lesions that block Pol ε, this slowing mechanism is likely a response to helicase-polymerase uncoupling (Sparks et al. 2019). We have shown that without priming, the CMG stalls at more stable sequences, supporting the “dead man’s switch” model. Furthermore, Pol ε remains with the CMG, and thus strand polymerization, and not the physical presence of a polymerase, must modulate helicase activity. In higher eukaryotes, an AT-rich unwinding element has not been found at origins, so we predict that Mcm2-7 will stall closer to origins in the absence of replication initiation in these systems. Our results suggest that in addition to impacting replisome progression, the mechanisms that regulate helicase-polymerase uncoupling contribute to replication initiation. Failure to regulate helicase uncoupling at the origin via a "dead man’s switch" may trigger replication catastrophe resulting from the generation of excess ssDNA and the sequestration of RPA (Toledo et al. 2013; Ercilla et al. 2020); thus mechanisms regulating helicase uncoupling from the replisome are likely conserved across eukaryotes and critical for preserving genome stability.

## Materials and Methods

### Yeast genetic methods

All yeast strains are in the W303 background and derived from HHY168 (Haruki et al 2008). Genotypes of all strains used are shown in Table 1. HHY168 and plasmids containing the FRB tag were obtained from Euroscarf. The LiAc/ssDNA/PEG method was used for all transformations, with the following adjustments (Gietz and Woods 2002). For all transformations involving temperature-sensitive strains or mutations, cells were grown at 24°C prior to transformation and an overnight incubation at 24°C was used in place of the typical 42°C heat shock. To amplify the FRB construct, primers for *CDC17* and *MRC1* were designed to match the 3’ end of each gene and the plasmids pFA6a-FRB-KanMX6 or pFA6a-FRB-HisMX6 (Longtine et al. 1998). These PCR products were then transformed into yeast. Temperature-sensitive mutations were added to CDC17 using CRISPR-Cas9 gene editing (Anand et al. 2017). First, guide RNA sequences were added to the Cas9-expressing plasmid bRA89, a gift from the Haber lab. To construct the repair template, a fragment of CDC17 was cloned into pCR 2.1 TOPO (Thermofisher #450641). The temperature-sensitive mutations, along with a silent mutation in the PAM sequence, were added to this fragment of CDC17 using site-directed mutagenesis (Agilent #200521). Both bRA89 with gRNA and the repair template (excised from the TOPO vector) were transformed into CDC17-FRB. RFA1 and POL2 were tagged with 3xFLAG using primers to the 3’ end of each gene and to pFA6a-6xGLY-3xFLAG-HIS3MX6 (Funakoshi and Hochstrasser 2009).

**Table 1.** Yeast strains used in this study.

| **Yeast strain** | **Genotype** | **Source & references** |
| --- | --- | --- |
| HHY168 | *Mata tor1-1 fpr1::NAT RPL13A-2 x FKB12::TRP1 leu2-3,112 can1-100 ura3-1 ade2-1 his3-11,15* | Haruki et al., 2008 |
| *CDC17-FRB* | *MATa, his3-11,15, BAR1::LEU, tor1-1, RPL13A-2xFKB12::TRP1, fpr1::NAT, CDC17-FRB::KanMX6, BrdU-Inc::URA3* | This paper |
| *cdc17-1-FRB* | *MATa, his3-11,15, BAR1::LEU, tor1-1, RPL13A-2xFKB12::TRP1, fpr1::NAT, cdc17-1-FRB::KanMX6, BrdU-Inc::URA3* | This paper. G904D mutation (Lucchini et al. 1990; Hartwell et al. 1973). |
| *cdc17-2-FRB* | *MATa, his3-11,15, BAR1::LEU, tor1-1, RPL13A-2xFKB12::TRP1, fpr1::NAT, cdc17-2-FRB::KanMX6, BrdU-Inc::URA3* | This paper, G637D mutation (Lucchini et al. 1990; Hartwell et al. 1973). |
| *cdc17-1,2-FRB* | *MATa, his3-11,15, BAR1::LEU, tor1-1, RPL13A-2xFKB12::TRP1, fpr1::NAT, cdc17-1,2-FRB::KanMX6, BrdU-Inc::URA3* | This paper. |
| *cdc17-1,2-FRB RFA1-3xFLAG* | *MATa, cdc17-1,2-FRB::KanMx6, RFA1-6xGLY-3xFLAG-HIS3MX6, BAR1::LEU, tor1-1, RPL13A-2xFKB12::TRP1, fpr1::NAT, BrdU-Inc::URA3* | This paper. |
| *cdc17-1,2-FRB POL2-3xFLAG* | *MATa, cdc17-1,2-FRB::KanMx6, POL2-6xGLY-3xFLAG-HIS3MX6, BAR1::LEU, tor1-1, RPL13A-2xFKB12::TRP1, fpr1::NAT, BrdU-Inc::URA3* | This paper. |
| *cdc17-1,2-FRB MRC1-FRB* | *MATa, cdc17-1,2-FRB::KanMX6, MRC1-FRB::HisMX6, BAR1::LEU, tor1-1, RPL13A-2xFKB12::TRP1, fpr1::NAT, BrdU-Inc::URA3* | This paper. |

### Cell synchronization and early S-phase arrest

All experiments were done in YPD media at 24°C unless indicated otherwise. For each experiment, cells were grown to saturation overnight, and this culture was used to inoculate a larger culture the next morning. Cells were doubled (based on OD_600_ readings) and then arrested for 2 hours using α-factor (Genway) at a final concentration of 50 ng/mL. In the final 30 minutes of the α-factor arrest, cultures were split in two and moved into different conditions: 24°C and 37°C alone or with rapamycin (1 mg/mL). At the end of the α-factor arrest, cells were washed twice with sterile water, resuspended in YPD media, and returned to their previous conditions (24°C or 37°C +/− rapamycin) for at least 1 hour.

### Spot assay

Cells were grown to saturation overnight in liquid culture and then all cultures were adjusted to the same OD_600_. Cells were serially diluted in ten-fold increments, mixed thoroughly, and spotted onto either YPD or YPD + rapamycin (1 mg/mL) plates. Plates were incubated 2 days at either 24°C or 37°C and then imaged.

### Flow cytometry

Experiments to analyze DNA content were performed as previously described (Gutiérrez et al. 2019).

### Chromatin preparation

25 mL samples were collected from cultures grown to a OD_600_ in the range of 0.8-1, cross-linked and quenched as previously described (Belsky et al. 2015). Cells were pelleted, flash-frozen, and stored at −80°C. MNase digestions were performed on thawed pellets again according to Belsky et al 2015. 1 ug of MNased DNA was used in the Illumina library preparations.

### Chromatin immunoprecipitation

ChIP experiments were performed as in Belsky et al 2015, with the following adjustments. 50 mL samples were collected from cultures grown to an OD_600_ of 0.8-1. For Mcm2-7 immunoprecipitation, 25 uL of AS1.1 antibody was used (Chen et al. 2007). For Rfa1-3xFLAG and Pol2-3xFLAG, 1-2 uL of mouse anti-FLAG (M2, F3165 Sigma) antibody was used. The entire ChIP pellet was used in the Illumina library preparations.

### Western blotting

1 mL samples were collected from cultures grown to OD_600_ in the range of 0.8-1, and prepared for analysis using TCA precipitation. Samples were run on a 10% SDS-PAGE gel and transferred to a nitrocellulose membrane with a wet transfer system. Membranes were blocked with 5% milk/1% Tween/TBS for 1 hour, then incubated overnight with goat polyclonal anti-Rad53 (Santa Cruz sc-6749, gift from S. Haase, Duke University) in the blocking solution. Membranes were washed and incubated in donkey anti-goat IgG-HRP (Santa Cruz sc-2020, gift from S. Haase) in blocking solution for 40 minutes at room temperature. Membranes were washed, incubated with ECL solution (Pierce #34095), and imaged using film.

### BrdU immunoprecipitation

5-bromo-deoxyuridine (BrdU) was added to cells at a final concentration of 400 ug/mL. At the end of the experiment, cells were pelleted, flash-frozen, and stored at −80°C. Genomic DNA was isolated from thawed cells and BrdU immunoprecipitation was performed as previously described (Ryba et al. 2011). The entire pellet resulting from the IP was used in the Illumina library preparations.

### Sequencing library preparation

MNase sequencing libraries were prepared as previously described (Henikoff et al. 2011; Belsky et al. 2015), and ChIP and BrdU IP libraries were prepared according to Illumina TruSeq protocols. The only adjustment made to both protocols was the use of the NEBNext Multiplex Oligos for Illumina kit (E7335S) in the adaptor ligation and PCR steps.

## Data analysis

Sequencing data was aligned to sacCer3 using Bowtie version 1.1.1 (Langmead et al. 2009). Data analysis was performed in R version 3.2.0 (R Core Team 2015). Scripts are available at https://gitlab.oit.duke.edu/rah60/hoffman-2021-paper. See supplemental methods for details on data analysis.

## Data access

All sequencing data from this study is available in the NCBI Sequence Read Archive (SRA) under accession number PRJNA704995.

## Acknowledgments

We thank current and former members of the MacAlpine laboratory and the Hartemink group for critical comments and suggestions. The antibodies for the Rad53 immunoblot were gifts from Steve Haase. This work was supported by the National Institutes of Health grant R35-GM127062 to D.M.M.

