## Supplemental Materials for "Disruption of origin chromatin structure by helicase activation in the absence of DNA replication"

### Supplemental Methods

*Genome & origin annotations*

An established list of 269 origins with an ORC footprint in both G1 & G2 phase of the cell cycle was used for all analyses (Belsky et al 2015). This set was narrowed to 239 origins by excluding origins close to chromosome ends or in regions with low coverage. Origin timing and efficiency annotations were from Belsky et al 2015 as well, with efficiency adapted from McGuffee et al 2013. Gene annotations were obtained from the Saccharomyces Genome Database (Cherry et al. 2012), and transcription factor binding sites were from (MacIsaac et al. 2006).

*MNase chromatin profiles and copy number*

All MNase chromatin profiling data was aligned using paired end mode in Bowtie (Langmead et al 2009). After determining that biological replicates from the MNase chromatin profiling experiments were highly correlated (Fig. S6A), replicates were merged. The resulting BAM files were all sampled to approximately the same number of reads for normalization. To visualize chromatin profiles, dot plots were constructed by obtaining fragment length and genomic position of all reads in a window around each origin, using the GenomicRanges package in R (Lawrence et al. 2013). A 2D density kernel estimate was used to calculate density values for each point in the window, and these density values were used to shade each point. Heatmaps of chromatin data were plotted by constructing matrices with rows corresponding to each origin and columns representing 100 bp bins of fragment counts around the origin. To plot copy number across a chromosome, coverage was obtained in 1 Kb bins for 140-180 bp reads in the MNase data. Copy number was then calculated by identifying a region with static coverage across all samples and dividing by mean coverage in this region. Data was smoothed using a moving average.

*Entropy score*

To calculate the entropy score for each origin, a matrix with a row for 10 bp increments of fragment lengths (20-250 bp) and columns for 10 bp bins around the origin +/- 1 Kb was constructed. The midpoint of each read was determined, and the matrix was populated with midpoint counts based on their position and original fragment length. The entropy function from the package 'entropy' was then used to calculate the 2D Shannon entropy estimate in bits from this matrix (Hausser and Strimmer 2009). The entropy estimate was calculated for both the G1 phase chromatin profile and 40 minutes, 37°C + rapamycin profile for each origin. The entropy estimate from G1 was subtracted from the 40 minutes entropy estimate to obtain the entropy score. This score reflects the change in entropy at 40 minutes from the G1 chromatin state. Distribution of the entropy score (Fig. 3A) was plotted using ggplot2 in R (Wickham 2016).

*BrdU aggregate origin plot*

Fragment counts in 1 bp bins were obtained in a 2 Kb window on either side of all origins. These counts were summed to obtain an aggregate BrdU profile around origins. Total counts within the window for each sample were scaled to the same total count. The profile was smoothed with a moving average and plotted. The replicates of BrdU-IP-seq data from WT cells released from G1 into hydroxyurea for 2 hours are available in the NCBI Sequence Read Archive (SRA) under accession number SRS597263 (Belsky et al 2015).

*ChIP-seq data*

Mcm2-7 ChIP-seq data from *cdc17-1,2*-FRB were aligned with single-end mode, while Mcm2-7 ChIP-seq data from *cdc17-1,2*-FRB MRC1-FRB was aligned in paired-end mode. All Rfa1 and Pol2 ChIP-seq data were aligned in paired end mode. We observed highly similar enrichment patterns at origins between the 40 and 60 minute timepoints in the Mcm2-7 ChIP-seq data, as well as between ChIP-seq data sets for Rfa1, Pol2, and Mcm2-7, demonstrating consistency across our datasets (Figures S6B & 4). BAM files were sampled to approximately the same number of reads for normalization purposes. Heatmaps of ChIP-seq data were constructed similarly to the chromatin heatmaps, except using 50 bp bins for genomic position. Matrices for these heatmaps were further normalized by scaling all matrices to contain the same total number of reads. For aggregate Mcm2-7 ChIP signal, a similar matrix was constructed and the sum of its columns calculated to obtain an aggregate Mcm2-7 signal around origins.

*Mcm2-7 peak positions*

Mcm2-7 peak positions in G1 phase and at 40 minutes (37°C + rapamycin) were obtained by calling peaks with MACS2, version 2.1.0.20151222 (Zhang et al. 2008). The command macs2 callpeaks was used with the --call-summits option to identify subpeaks. Matching input datasets were used as controls for each ChIP data set. The best G1 peak at each origin was determined by identifying all peaks +/-500 bp from the origin center and selecting the peak within this set with the highest enrichment value as determined by MACS2. The top peaks at each origin at 40 minutes were determined by identifying all peaks +/-2000 bp from the origin center and excluding any peaks within

100 bp of the previously identified G1 peak position. The most enriched peak upstream and downstream of the origin was then identified from the remaining peaks.

*Sequence analysis*

The BSGenome package in R was used to obtain sequences with a given genomic range (Pagès 2015). The number of each nucleotide present was counted to determine GC or AT content of a given sequence. The length of the AT-rich region at each origin was determined by stepping out from the annotated origin center in 50 bp bins until GC content exceeded 0.4. This point was recorded as a boundary of the AT region, and the stepping process was repeated on the other side of the origin. The locations of the boundaries on either side of the origin were used to calculate the center and length of the AT-rich region which was used to align and order origins in Fig. 6. For the AT content heatmap, a matrix with a row for each origin and columns containing “0” for 3 bp bins was created. 3-mers containing only A, T or both were counted, and the corresponding bin set to “1.” The values in this matrix were used to shade the heatmap.

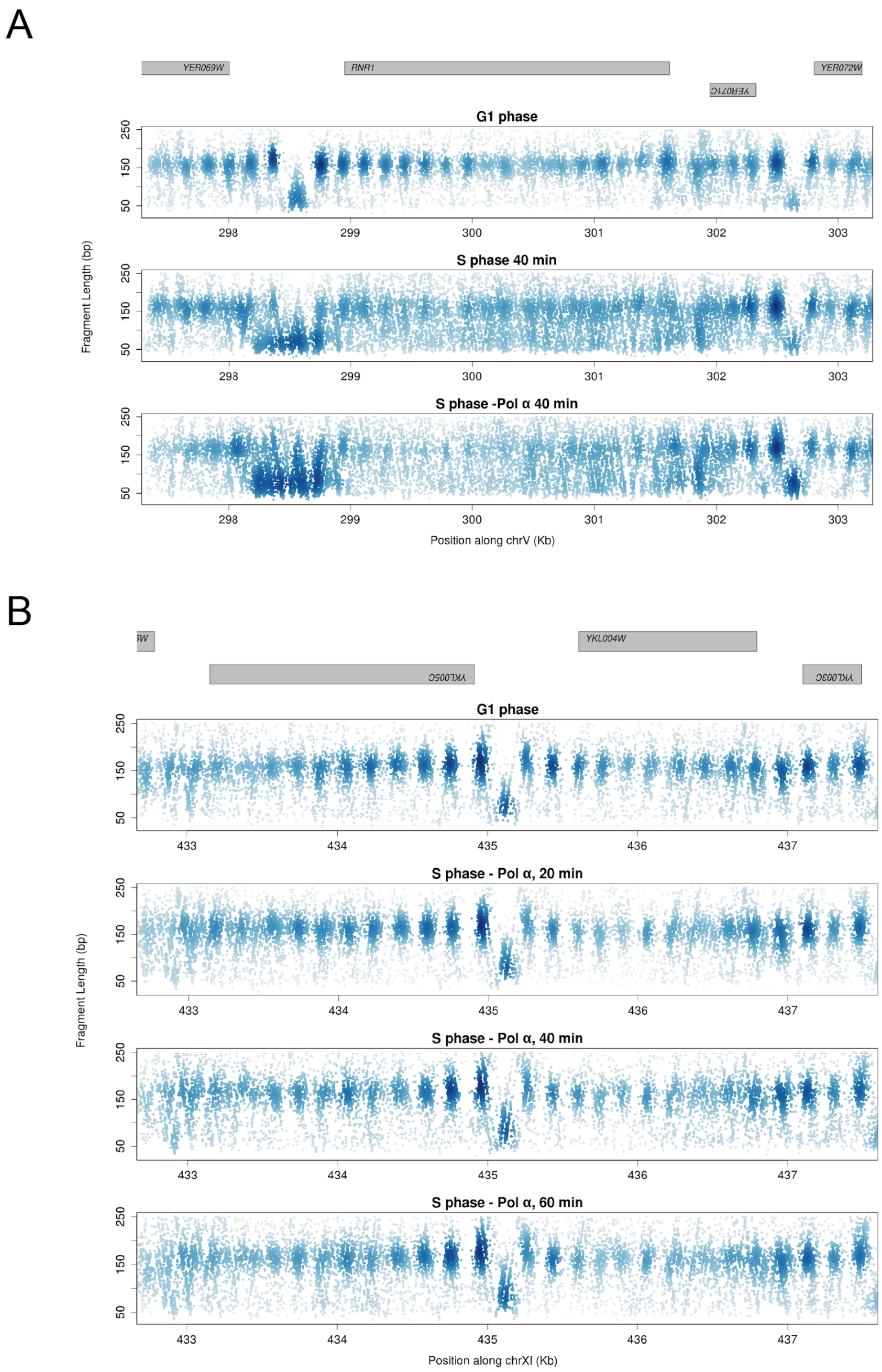


**Figure S1**: MNase chromatin profiles in *cdc17-1,2-FRB* cells at other genomic loci. (**A**) Chromatin profile for *RNR1*, a gene expressed at the G1/S phase transition, in G1 phase and released into S phase at either 24°C or 37°C +rapamycin. Gene bodies are indicated in gray at the top of the plot. (**B**) Chromatin profile for an Abf1 site on Chromosome XI at 37°C + rapamycin in *cdc17-1,2-FRB* cells. Cells were synchronized in G1 phase and then released into S phase under restrictive conditions.

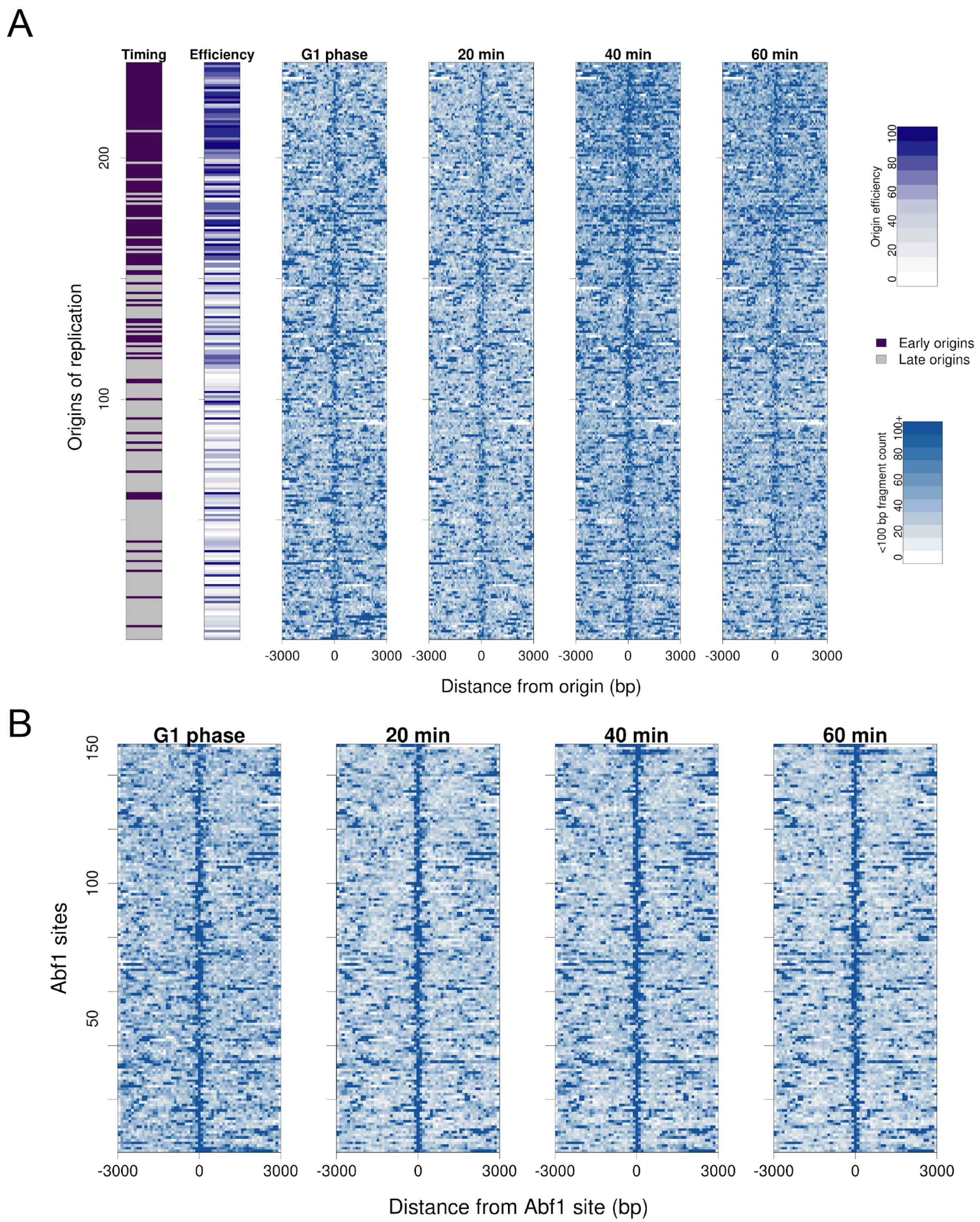


**Figure S2**: Chromatin disorganization does not occur in *cdc17-1,2-FRB* cells at origins under permissive conditions or at Abf1 sites under restrictive conditions. (**A**) Heatmap of 239 replication origins in *cdc17-1,2-FRB* cells at 24°C, in G1 and released into S phase. Origins are ordered as in **Figure 3B**. The midpoints of <100 bp fragments resulting from MNase digestion were counted around each origin, and are plotted in 100 bp bins. Origin timing and efficiency deciles are annotated on the left. Time labels reflect minutes since release from G1 phase. (**B**) Heatmap of 151 Abf1 binding sites at 37°C + rapamycin in *cdc17-1,2-FRB* cells, at G1 and released into S phase. Abf1 sites are ordered by decreasing entropy score, reflecting the difference between 40 min, 37°C + rapamycin entropy and G1 phase entropy at Abf1 sites. The midpoints of <100 bp fragments resulting from MNase digestion were plotted as in **S2A**.

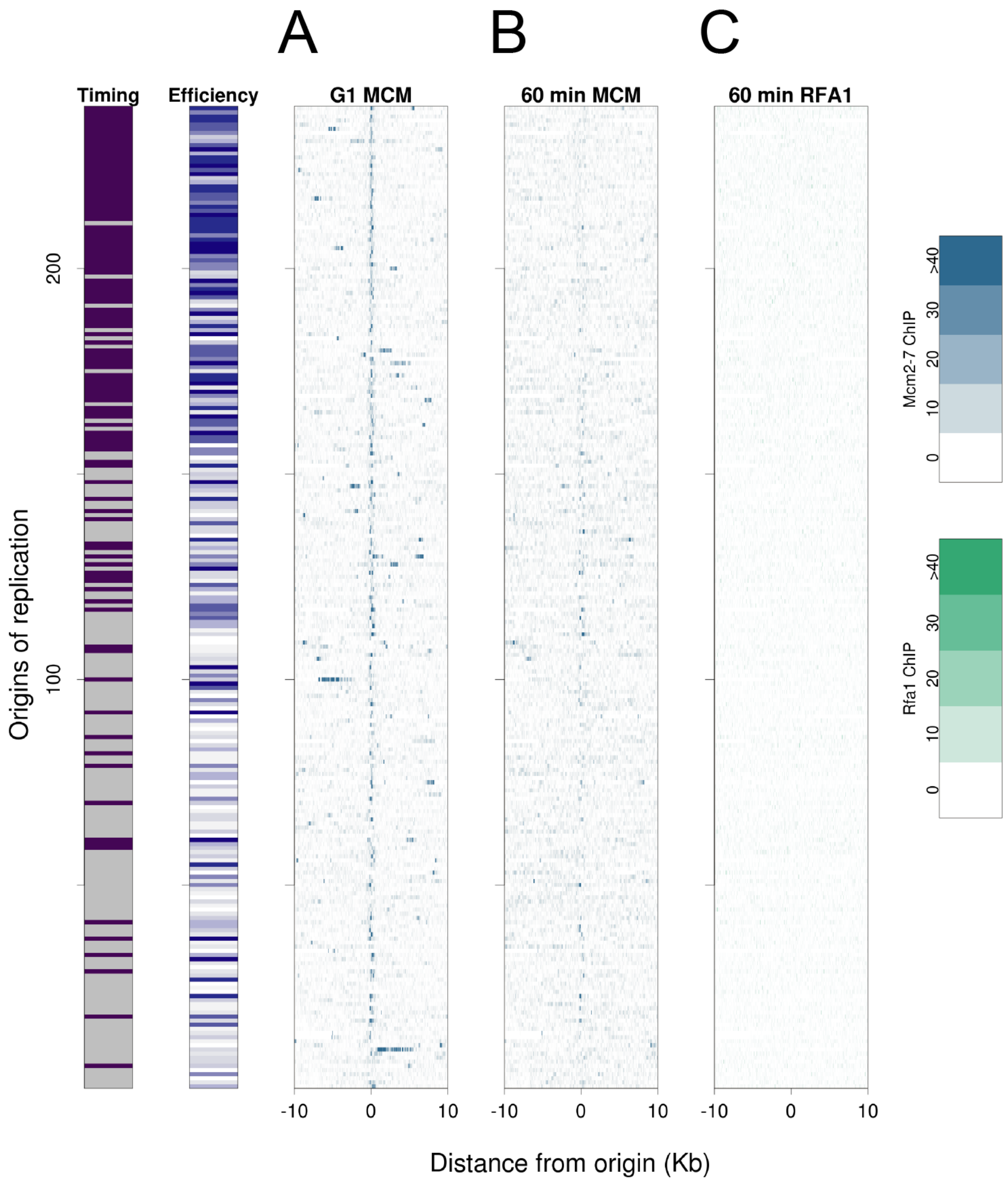


**Figure S3**: Mcm2-7 and Rfa1 are not highly enriched around origins at 24°C in S phase in *cdc17-1,2-FRB* cells. Origins are ordered as in **Figure 3B**. (**A**) Mcm2-7 ChIP-seq enrichment around origins in G1 phase and (**B**) 60 minutes into S phase at 24°C. (**C**) Rfa1 ChIP-seq enrichment around origins 60 minutes into S phase at 24°C. G1 Rfa1 ChIP enrichment is subtracted from the 60 minute enrichment.

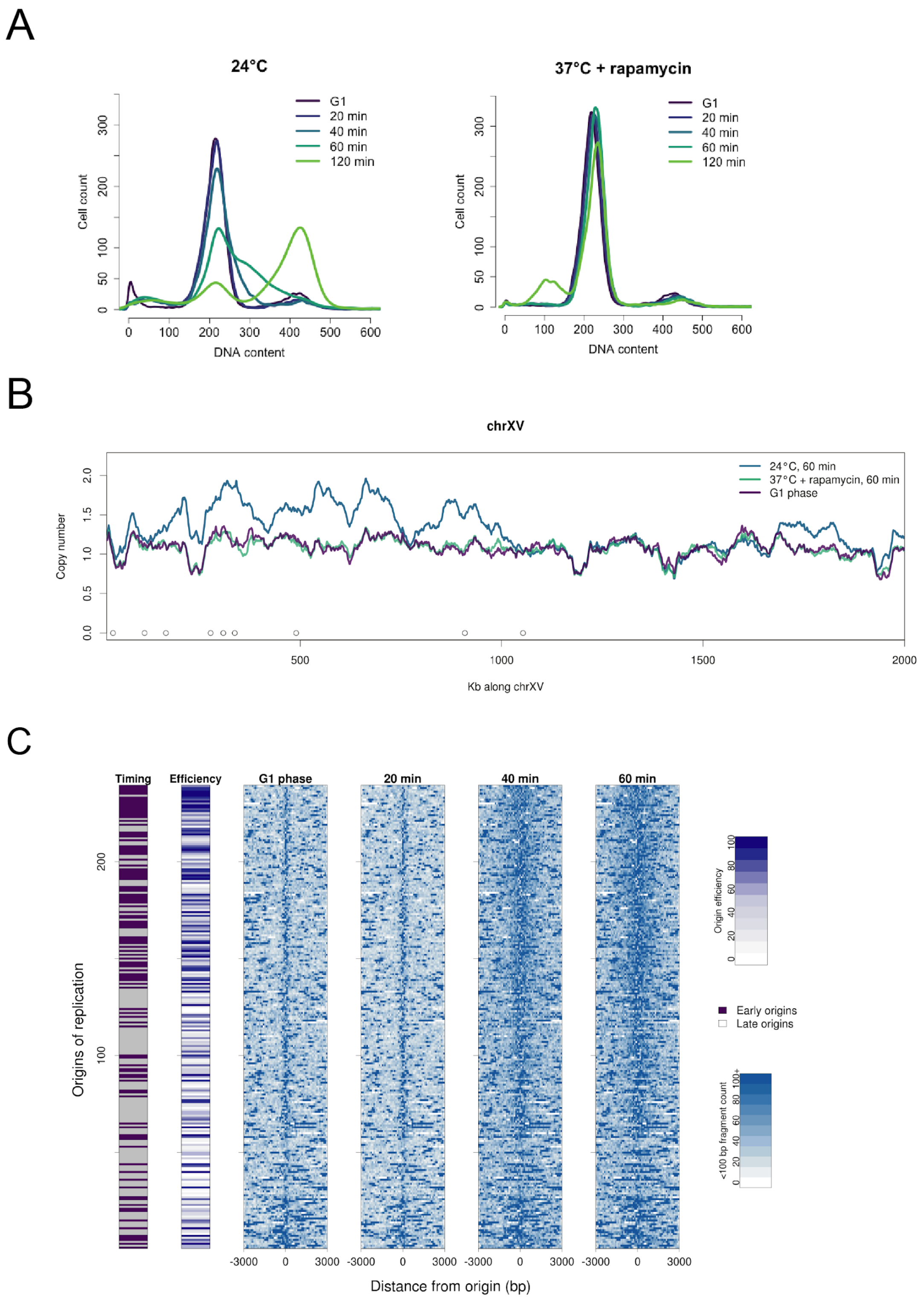


**Figure S4**: *cdc17-1,2-FRB MRC1-FRB* cells arrest prior to replication and have disorganized chromatin around early and late origins under restrictive conditions. (**A**) DNA content measured by flow cytometry in *cdc17-1,2-FRB MRC1-FRB* cells at restrictive (37°C + rapamycin) and permissive conditions (24°C) during G1 phase and up to 120 minutes after release from G1. Experiment performed as shown in **Figure 1B**. (**B**) Copy number across Chromosome XV during G1 or 60 minutes after release from G1, under permissive or restrictive conditions. Open circles along the x-axis denote the positions of early origins. (**C**) Heatmap of 239 replication origins under restrictive conditions in *cdc17-1,2-FRB MRC1-FRB* cells, ordered by decreasing entropy score, reflecting the difference between G1 phase entropy and 40 min, 37°C + rapamycin entropy. The midpoints of <100 bp fragments resulting from MNase digestion were counted around each origin, and are plotted in 100 bp bins. Origin timing and efficiency deciles are annotated on the left. Time labels reflect minutes following release from G1 phase.

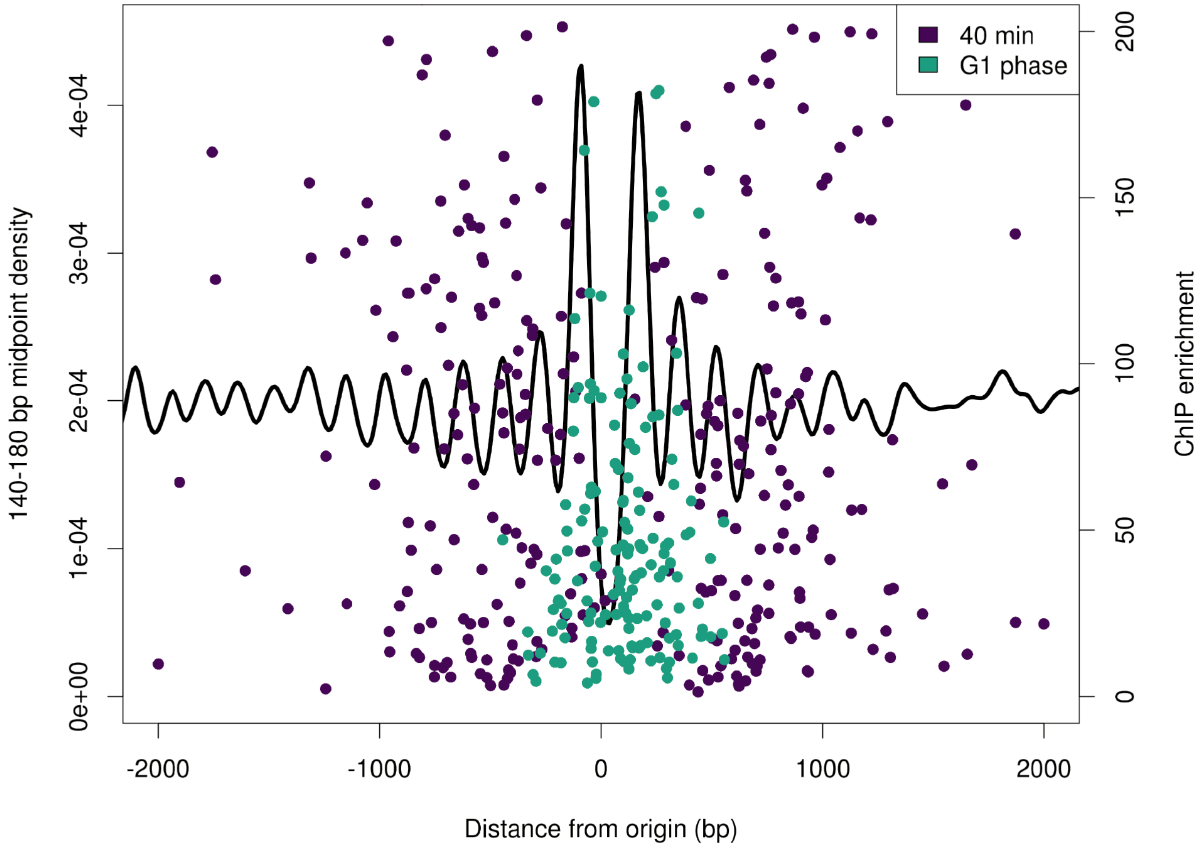


**Figure S5**: Mcm2-7 ChIP-seq peaks in *cdc17-1,2-FRB* cells do not align with specific chromatin features around origins. Density of nucleosome-sized fragments (between 140-180 bp) in G1 phase at 239 replication origins is plotted alongside Mcm2-7 ChIP-seq peak positions from G1 phase and 40 minutes after release from G1 phase at restrictive conditions. Enrichment value was determined during peak calling with MACS2.

A

| Sample name | $R^2$ | Strain |
|---|---|---|
| G1 phase 37°C + rapamycin | 0.935 | cdc17-1,2-FRB |
| 20 min 24°C | 0.973 | cdc17-1,2-FRB |
| 40 min 24°C | 0.951 | cdc17-1,2-FRB |
| 60 min 24°C | 0.955 | cdc17-1,2-FRB |
| 20 min 37°C + rapamycin | 0.943 | cdc17-1,2-FRB |
| 40 min 37°C + rapamycin | 0.924 | cdc17-1,2-FRB |
| 60 min 37°C + rapamycin | 0.969 | cdc17-1,2-FRB |
| G1 phase 37°C + rapamycin | 0.932 | cdc17-1,2-FRB MRC1-FRB |
| 20 min 37°C + rapamycin | 0.979 | cdc17-1,2-FRB MRC1-FRB |
| 40 min 37°C + rapamycin | 0.981 | cdc17-1,2-FRB MRC1-FRB |
| 60 min 37°C + rapamycin | 0.976 | cdc17-1,2-FRB MRC1-FRB |

B

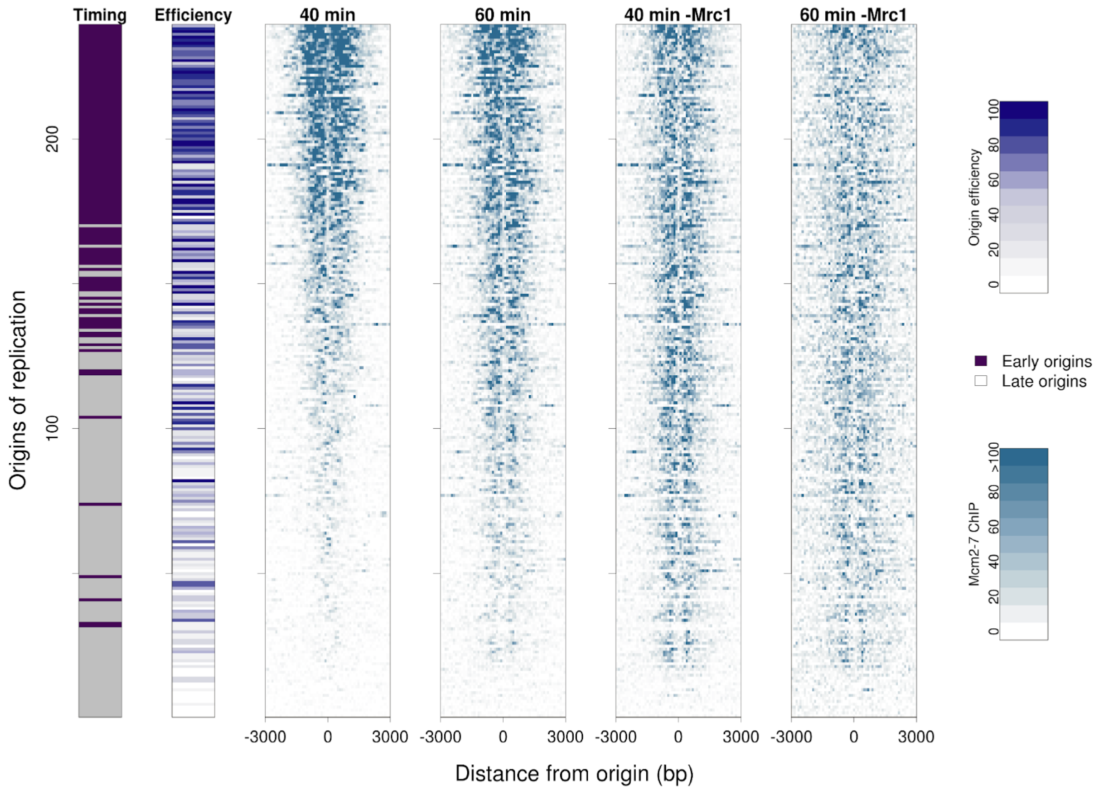


**Figure S6:** Chromatin and ChIP-seq data is highly consistent across replicates and between time points. (**A**) Correlation of coverage in MNase replicates across chromosome IV in *cdc17-1,2-FRB* and *cdc17-1,2-FRB MRC1-FRB* cells. Coverage was obtained in 1 Kb bins across Chromosome IV for each replicate, and then the coefficient of determination was calculated using a two-sided Pearson correlation. (**B**) Heatmap of Mcm2-7 ChIP-seq enrichment at 239 origins 40 and 60 minutes after release from G1 under restrictive conditions in *cdc17-1,2*-FRB and *cdc17-1,2*-FRB *MRC1-FRB* (noted as -Mrc1) cells. Origins are ordered by decreasing Mcm2-7 ChIP-seq enrichment in the "40 min" panel.
